# Divergent signaling profiles in mTOR gain-of-function Smith-Kingsmore syndrome (SKS) and TSC2 deficiency

**DOI:** 10.64898/2026.05.09.724025

**Authors:** Curtis R. Carlson, Yang Shen, Hongzhi He, Erwin K. Gudenschwager, Chunyan Hou, Junfeng Ma, Joanna C. Chiu, Andrew C. Liu

## Abstract

Smith-Kingsmore syndrome (SKS) is a rare neurodevelopmental disorder caused by gain-of-function mutations in *MTOR*, yet whether these mutations phenocopy *TSC2* loss or establish a distinct signaling state remains unclear. Using quantitative proteomics, phosphoproteomics, and transcriptomics in isogenic cell models of SKS (*MTOR^Δ4aa^*), *TSC2* loss (*TSC2*^−/–^), and wild-type controls under glucose depletion and refeeding, we find that *MTOR^Δ4aa^* and *TSC2*^−/–^ cells occupy fundamentally distinct regulatory states. *TSC2*^−/–^ cells exhibit broad anabolic remodeling and a transcriptional program dominated by NF-κB- and STAT-driven inflammatory responses. *MTOR^Δ4aa^* cells instead display enrichment of nuclear and RNA processing programs, E2F/MYC-driven transcription, and a constrained proteomic dynamic range across nutrient states. Phosphoproteomic analysis of *MTOR^Δ4aa^* reveals rerouting of nutrient-responsive signaling toward MAPK/ERK- and Ca^2+^/CaMK-dependent pathways with limited canonical mTORC1/S6K1 engagement. These findings establish SKS as a signaling rewiring disorder distinct from classical mTORC1 hyperactivation, with implications for therapeutic targeting.

## Introduction

Smith-Kingsmore syndrome (SKS; MIM 616638) is a rare neurodevelopmental disorder caused by *de novo* gain-of-function mutations in the *MTOR* gene^1,2^. Individuals with SKS present with a broad spectrum of core clinical symptoms, including macrocephaly/megalencephaly, developmental delay, intellectual disability, autism spectrum disorder, seizures, and epilepsy^1,3,4^. More recent work has expanded the clinical manifestations to include hyperactivity, self-injurious behavior, hyperphagia, heat intolerance, and sleep-wake disturbances, indicating systemic homeostatic imbalance^4^. To date, over 30 distinct alleles have been reported in over 100 individuals worldwide, with the majority of mutations clustering in the FAT and kinase domains of mTOR^2,4^. Several SKS variants have been shown to exhibit increased mTOR activity, characterized by elevated phosphorylation of canonical targets of mTORC1 (S6K1, S6, and 4E-BP1)^4–6^ and mTORC2 (AKT)^5^, along with varying degrees of resistance to rapamycin-induced inhibition^4,6^. However, these canonical target readouts provide only a narrow view of pathway activity. The mechanisms by which distinct *MTOR* mutations associated with SKS alter nutrient sensing, signaling dynamics, and downstream cellular states, leading to clinical phenotypes, remain poorly understood. Therefore, understanding how SKS mutations impact mTOR signaling is essential for defining disease mechanisms and guiding therapeutic strategies for SKS.

mTOR is a serine/threonine kinase that acts as a central integrator of nutrients, growth factors, and cellular energy to regulate anabolic metabolism^7^. It is the catalytic subunit of two distinct multiprotein complexes, mTOR Complex 1 (mTORC1) and mTOR Complex 2 (mTORC2). mTORC1 is acutely sensitive to nutrient status, integrating glucose and growth factor signals via the small GTPase RHEB^8^ and amino acid signals through RAG GTPases at the lysosomal surface^9^. Under nutrient-replete conditions, mTORC1 promotes protein, lipid, and nucleotide synthesis while suppressing autophagy. mTORC2 primarily responds to growth factors to regulate cytoskeletal dynamics, cell survival, glucose uptake, and ion transport through activation of several AGC kinases, including AKT, SGK, and PKC^10^. Together, mTORC1 and mTORC2 coordinate cellular metabolic homeostasis, and their dysregulation underlies a spectrum of metabolic and neurodevelopmental disorders collectively termed “mTORopathies”^11^.

In the mTORC1 pathway, the tuberous sclerosis complex proteins TSC1 and TSC2 function as a GTPase activating protein (GAP) complex that inhibits RHEB and suppresses mTOR activity under conditions of limited growth factor or energy availability. Loss-of-function mutations in *TSC1* or *TSC2* cause tuberous sclerosis complex (TSC), an mTORopathy characterized by non-cancerous tumors, seizures, and intellectual disability^12^. At the molecular level, TSC1/2 loss leads to constitutive hyperactivation of mTORC1 while only indirectly affecting mTORC2 activity^13^. Rapamycin and its analogs (rapalogs) are clinically approved to treat TSC and can improve patient outcomes by suppressing mTORC1 activity. In contrast, SKS mutations directly affect the mTOR kinase itself, raising the possibility that SKS mutations affect both mTORC1 and mTORC2. Consistent with this, SKS variants show variable responses to rapamycin, requiring careful tuning of dosage^4,14^. These clinical and mechanistic differences suggest that SKS represents a distinct mode of mTOR network dysregulation, rather than a simple recapitulation of canonical mTORC1 hyperactivation.

To test whether SKS mutations phenocopy classical mTORC1 hyperactivation, as observed in TSC, or instead adopt distinct modes of signaling, we analyzed nutrient responses in isogenic human U2OS cell models: wild-type (WT) cells; *MTOR^(ΔR1480-C1483/ΔR1480-C1483)^* (hereinafter referred to as *MTOR^Δ4aa^*), an SKS knock-in allele previously shown to be hyperactivating^4^; and *TSC2* knockout cells (*TSC2*^−/–^) as a model of classical mTORC1 hyperactivation. Because mTORC1 activity is reduced under nutrient depletion and reactivated upon nutrient restoration^4,15^, we subjected all cell models to acute glucose depletion to assess basal mTOR activity, followed by refeeding to probe glucose-induced signaling dynamics. We applied parallel quantitative proteomic and phosphoproteomic analyses to capture both proteome remodeling and kinase signaling dynamics across glucose depletion and refeeding conditions. In addition, we performed RNA-seq under glucose-depleted conditions to define the basal transcriptional state associated with each genotype. This integrated analysis revealed that, unlike *TSC2*^−/–^ cells, *MTOR^Δ4aa^*cells exhibit non-canonical nutrient-responsive signaling, characterized by a distinct basal transcriptional and proteomic identity, a constrained proteomic dynamic range across glucose conditions, and a shift away from canonical mTORC1-driven anabolic signaling toward MAPK/ERK and Ca^2+^/CaMK pathways. Together, these findings indicate that the *MTOR^Δ4aa^* SKS variant establishes a signaling state not fully captured by classical mTORC1 hyperactivation.

## Materials and Methods

### Cell lines

The *MTOR^Δ(R1480–C1483)^* mutation was introduced into human U2OS cells by CRISPR gene editing to generate a knock-in gain-of-function allele, referred to here as *MTOR^Δ4aa^*. The generation and validation of these cells were described previously^4^.

To generate *TSC2* knockout (*TSC2^−/–^*) cell lines, CRISPR-Cas9-mediated genome editing was performed in U2OS cells. Two single-guide RNAs (sgRNAs) were designed to target the human *TSC2* locus flanking a large coding region of exons 2-12, resulting in a genomic deletion upon dual cutting. sgRNA target sequences were: sgRNA-1: 5′-TGGTGCGTCCTGGTCCACCA-3′; and sgRNA-2: 5′-CGTCCATGACCTGTTGACCA-3′. sgRNAs were selected based on predicted on-target efficiency and minimal off-target potential. sgRNA1 and sgRNA2 were cloned into pX458-GFP and pX458-mCherry, respectively, as described^16,17^. These CRISPR vectors were introduced into U2OS cells using standard transfection methods. Following transfection, cells were dissociated for 2-color FACS sorting. Sorted cells were pooled, followed by single-cell seeding on 96-well plates by limiting dilution and characterization of clonal cell lines.

Genomic DNA was extracted from parental cells, pooled CRISPR-edited populations, and individual clones. Deletions at the *TSC2* locus were screened by PCR using multiple primer combinations spanning the targeted region. Primer sequences used for genotyping were: a-F51: 5′-TCCTGCTTCACATGGGTAGAG-3′; b-R52: 5′-CCCAGCCTTCTCTGCATTT C-3′; c-F31: 5′-ATGTGGGGAGTGGAAGTCAG-3′; and d-R32: 5′- ACACAGGTGGGAGGAAACTG-3′. In wild-type cells, primer pairs a-F51/b-R52 and c-F31/d-R32 amplified fragments of 551 bp and 400 bp, respectively. In *TSC2^−/–^* cells, the a-F51/d-R32 primer pair produced a 662 bp fragment (instead of a large 13,965 bp product), distinguishing deletion alleles from wild-type. Selected clones were validated by Sanger sequencing to confirm precise junctions of the deletion. Both homozygous and heterozygous deletion clones were identified and used for downstream analyses.

### Cell culture

mTOR signaling was assessed under glucose depletion (basal) and restimulation with complete growth medium (refeeding) as previously described^4^. Basal medium consisted of D-PBS, 20 mM HEPES, and 15 mg/L phenol red. The glucose-free medium was prepared by supplementing this basal medium with amino acids and dialyzed fetal bovine serum (Gibco A33820-01; 10-kDa cutoff), which contains low glucose (<5 mg/dL) and amino acids. Cells were grown in standard growth medium, then washed and incubated in glucose-free medium for 1 or 2 h (“Depleted”) to establish a basal mTOR activity state. For glucose restimulation (“Fed”), cells were returned to regular high-glucose growth medium for 1 or 2 h. Following treatment, cells were rapidly harvested on ice. A subset of samples was processed for global proteomics and phosphoproteomics to quantify mTOR-dependent signaling and phosphorylation responses across depletion and refeeding conditions. Parallel samples from selected conditions were collected for RNA extraction and downstream RNA-seq analysis to assess transcriptional responses.

### mTOR activity assay by immunoblotting

mTOR activity in *MTOR^Δ4aa^* and *TSC2^−/–^* cells was assessed by immunoblotting for canonical downstream targets. Briefly, cell lysates were analyzed for phosphorylation of ribosomal protein S6 (p-S6) with total S6 and β-actin used as loading controls. Rapamycin treatment was used where indicated. Experimental procedures, antibodies, and quantification methods were performed as described previously for mTOR activity assays^4^.

### Cellular circadian rhythm analysis

To examine the impact of *TSC2* loss on cellular circadian properties, wild-type and *TSC2^−/–^* U2OS cells expressing the circadian luciferase *Per2-dLuc* reporter rhythm was recorded on Lumicycle, a real-time bioluminescence recording device. Raw luminescence data (counts/second) as a function of time (days) were analyzed with the LumiCycle Analysis program (version 2.53, Actimetrics) to determine circadian parameters, as described previously^4^.

### Mass spectrometry sample preparation and NanoUPLC-MS/MS

Total proteins were extracted from cell pellets for quantitative proteomics, by using a similar procedure published previously^18^. In brief, equal amount of proteins from each sample (n = 3 biological replicates per condition) (300 µg) was suspended with 50 µL of 5% SDS buffer (containing 50 mM TEABC and 20 mM DTT) and heated for 10 min at 95°C. After cooling down to room temperature, 4 µL of 500 mM IAA in 5% SDS solution was added to a final concentration of 40 mM and incubated in the dark for 30 min. Undissolved matter was centrifuged for 8 min at 13,000 x g. The supernatant was saved and used for downstream processing using a S-Trap column (ProtiFi, LLC, Fairport, NY). Proteins were digested with sequencing-grade Lys-C/trypsin (Promega) by incubation at 37°C overnight. An aliquot was saved for total proteomics, and the remaining peptides (∼95%) were used for phosphopeptide enrichment by using a High Select™ Phosphopeptide Enrichment Kit (Thermo Fisher Scientific, Waltham, MA), by following the manufacturer’s manual. The resulting peptides were eluted and dried down with a SpeedVac (Fisher Scientific, Pittsburgh, PA).

Peptides were analyzed with a reversed phase nanoElute 2 (Bruker Daltonics Gmbh., Bremen, DE) coupled with timsTOF Ultra mass spectrometer (Bruker Daltonics Gmbh). Samples in 0.1% FA solution were loaded and separated with an analytical column (PepSep C18 packed with ReproSil AQ as stationary phase, pore size 120 Å, 10 cm x 75 µm, 1.9 µm) with the temperature maintained at 50°C. The flow rate was set at 300 nL/min. A 40-min gradient of buffer A (0.1% formic acid) and buffer B (0.1% formic acid in ACN) was used for separation: 2% buffer B at 0 min, 23% buffer B at 28 min, 30% buffer B at 30 min, 90% buffer B at 36 min, and 90% buffer B at 40 min. The CaptiveSpray source was used, with the ion spray voltage 1600 V. All data were acquired by the timsTOF Ultra mass spectrometer via Bruker timsControl in diaPASEF mode. Full MS data were acquired in a range of 100–1700 m/z and 1.45 1/K0 [V-s/cm2] to 0.64 1/K0 [V-s/cm2] in diaPASEF. DIA windows ranged from 400 m/z to 1000 m/z and were acquired with ramp times of 100 ms or otherwise indicated. High sensitivity detection for low sample amounts was enabled without diaPASEF data denoising.

### Data processing, quantitative proteomics, and phosphoproteomics analysis

Analysis of DIA raw files were conducted by using Spectronaut 17 (Biognosys, Schlieren, CH) in the directDIA mode. All settings were default. In brief, dynamic retention time prediction with local regression calibration was selected. Interference correction on MS and MS2 level was enabled. The Qvalue Cutoff was set to 1% at peptide precursor and protein level using scrambled decoy generation and dynamic size at 0.1 fraction of library size. MS2-based quantification by area was used, enabling local cross-run normalization.

Raw phosphopeptide intensities with localization scores >0.75 were exported from Spectronaut and aggregated to the phosphosite level using a custom Python script “sum_phospho_sites.py”. Intensity values for all peptides mapping to the same modified residue were summed to generate single phospho-site level intensities.

Raw total protein intensities were exported from Spectronaut and imported into DEP2^43,44^ (v 0.3.7.3)^19^ and normalized using VSN normalization. Missing protein values were imputed using the “mixed on proteins” setting. Pairwise comparisons between genotypes and nutritional status were performed on the normalized and imputed datasets. Resulting *p*-values were adjusted for multiple hypothesis testing using the Benjamini-Hochberg correction to control false discovery rate (FDR). Proteins with an adjusted *p*-value ≤ 0.05 were considered differentially expressed.

Raw aggregated phosphosite intensities were also imported in DEP2 (v 0.3.7.3) for normalization and imputation using the same settings as above. Phosphosite intensities were further normalized to total protein levels using the “protein quantification correction” step in the DEP2 package. Pairwise comparisons between genotypes and nutritional status were performed on the normalized and imputed datasets and the resulting *p*-values were adjusted for multiple hypothesis testing using the Benjamini-Hochberg correction to control FDR. Phosphosites with an adjusted *p*-value ≤ 0.05 were considered differentially expressed.

Enrichment analysis of differentially expressed proteins was conducted using the WebGestalt^20,21^ online portal (last accessed May 16^th^, 2025). For Gene Set Enrichment Analysis (GSEA), proteins were ranked by Log_2_ Ratio and mapped to either the MSigDB Hallmark Gene Set database^22^, the WikiPathways database^23^, or the CORUM protein complex database^24,25^. Significance level of enriched GSEA categories was determined using the “TOP” approach implemented in WebGestalt, where categories are first ranked by FDR and then the top 10 positively or negatively enriched categories are selected. Outputs were further pruned for redundant protein mapping by using the “weighted set cover” option in WebGestalt.

For Over-Representation Analysis (ORA), proteins from each pairwise comparison were first filtered by adjusted *p*-value ≤ 0.05. Differentially expressed proteins were then compared between pairwise contrasts (e.g. *MTOR^Δ4aa^*fed vs WT fed against *TSC2*^−/–^ fed vs WT fed) to identify shared and unique hits across conditions. These shared and unique protein subsets were then input into WebGestalt for ORA using the Gene Ontology (GO) Biological Processes (BP) database^26,27^ using the same settings as above.

For kinome enrichment analysis, each identified phosphosite was converted into a centered 21-residue sequence window by appending up to ± 10 flanking amino acids from the parent protein, positioning the phosphorylated residue at the center using the custom Python script “center_peptides.py”. Kinome enrichment was performed using Fisher Enrichment Analysis as implemented in The Kinase Library^28,29^ web portal (v. 1.2.0, last accessed April 10^th^, 2025). Centered phosphosites were imported and ranked by Log_2_ Ratio. Enrichment was determined by using the “Percentile Rank” metric, with significantly activated/inhibited kinases defined using an enrichment threshold of 15 and adjusted *p*-value ≤ 0.1 for serine/threonine kinases, and an enrichment threshold of 8 and *p*-value ≤ 0.05 for tyrosine kinases. Data visualization was performed using the ggplot2 package in R (v.4.2.2). Note that kinase enrichment scores reflect motif-based predictions of likely kinase activity from substrate phosphorylation patterns.

### RNA-seq library preparation, sequencing, and analysis

Total RNA was extracted from cultured cells (n = 3 biological replicates per condition) using TRIzol reagent (Thermo Fisher Scientific) followed by chloroform phase separation and column purification with the RNeasy Mini kit (Qiagen) according to the manufacturers’ protocols. RNA samples were treated with DNase I to remove residual genomic DNA. RNA concentration and purity were assessed using a NanoDrop spectrophotometer (Thermo Fisher Scientific), and RNA integrity was evaluated using an Agilent 2100 Bioanalyzer. All samples exhibited RNA integrity numbers (RIN) greater than 8.0 and were used for library preparation. Poly(A)-selected RNA-seq libraries were generated using the Illumina mRNA Prep kit. Libraries were sequenced on an Illumina NovaSeq platform using paired-end 2 × 100 bp reads to a depth of at least 40 million reads per sample. Sequencing was performed at the University of Florida ICBR NextGen DNA sequencing core facility.

Raw sequencing reads (FASTQ files) were processed on the University of Florida HiPerGator high-performance computing cluster. Reads were aligned to the human reference genome (GRCh38) using HISAT2 (v2.2.1). Gene-level read counts were generated using featureCounts (v2.0.1) based on the GENCODE gene annotation (GRCh38).

Raw gene counts were imported and processed using DESeq2^30^ implemented in DEP2 (v 0.3.7.3) using default settings. Genes with low counts across all samples were filtered prior to differential expression analysis. Differential expression between conditions was assessed using the Wald test implemented in DESeq2, and p-values were adjusted for multiple hypothesis testing using the Benjamini–Hochberg procedure. Genes with an adjusted *p*-value ≤ 0.05 were considered differentially expressed. Transcription factor activity enrichment was determined using the R packages dorothea (v 1.10.0)^31^ and decoupleR (v 2.4.0) in R (v.4.2.2)^32^. Transcription factor activity scores were calculated using the univariate linear model (ULM) method based on expression changes of curated target gene sets.

## Results

### Isogenic Cell models to interrogate SKS-associated mTOR signaling

Smith-Kingsmore syndrome (SKS) arises from gain-of-function mutations in *MTOR* that result in aberrant mTOR signaling^3,4^. To test whether an SKS mutation phenocopies classical mTORC1 hyperactivation or instead adopts a distinct nutrient-sensing state, we generated an isogenic human U2OS cell panel comprising wild-type (WT), an mTOR gain-of-function SKS variant *MTOR^Δ4aa^*, and *TSC2*^−/–^ cells as a benchmark for classical mTORC1 hyperactivation^33^. In a previous study, the *MTOR^Δ4aa^* mutation was introduced into U2OS cells by CRISPR knock-in to generate an endogenous gain-of-function allele^4^. The mutant cells displayed elevated basal signaling consistent with pathogenic mTOR activation observed in SKS. These *MTOR^Δ4aa^* cells were used in the current study for multi-omics profiling.

To establish *TSC2*^−/–^ cells as a model of constitutive mTORC1 activation, we generated knockout clones using CRISPR-Cas9-mediated deletion of a ∼13.9 kb genomic region spanning exons 2-12 of the *TSC2* locus (Figure S1A). PCR genotyping and Sanger sequencing confirmed successful deletion of the targeted region, and immunoblot analysis verified loss of TSC2 protein in homozygous clones (Figure S1B-C). Consistent with the role of the TSC1/2 complex as a negative regulator of mTORC1, *TSC2*^−/–^ cells displayed elevated basal phosphorylation of ribosomal protein S6 (p-S6 Ser235/236), indicating constitutive mTORC1 signaling (Figure S1D). In addition, *TSC2*^−/–^ cells exhibited a shortened circadian period and reduced rhythm amplitude in the *Per2-dLuc* bioluminescence reporter assay, phenotypes previously associated with mTOR hyperactivation (Figure S1E-F)^4,34–36^.

We next compared mTOR signaling activity across WT, *MTOR^Δ4aa^*, and *TSC2*^−/–^ cells under defined nutrient conditions. Each genotype was subjected to acute glucose depletion followed by refeeding, conditions that suppress glucose-dependent mTOR signaling toward a basal state and subsequently re-engage nutrient-responsive mTOR activity, respectively, allowing us to probe how SKS alters these dynamics^37,38^. More specifically, cells underwent 1 hour of glucose depletion to establish a low-nutrient baseline, followed by 1 hour of glucose restimulation to acutely activate mTOR (Figure 1A). Under depleted conditions, basal mTOR signaling exhibited a clear genotype-dependent hierarchy: *TSC2*^−/–^ cells showed the highest levels of p-S6, followed by *MTOR^Δ4aa^* cells, and then WT controls (Figure 1B, compare lanes 5, 3, and 1), consistent with gain-of-function signaling states in the mutant cells. Glucose restimulation increased mTOR signaling in all genotypes. These matched samples were subsequently subjected to quantitative proteomic and phosphoproteomic profiling.

**Figure 1.**
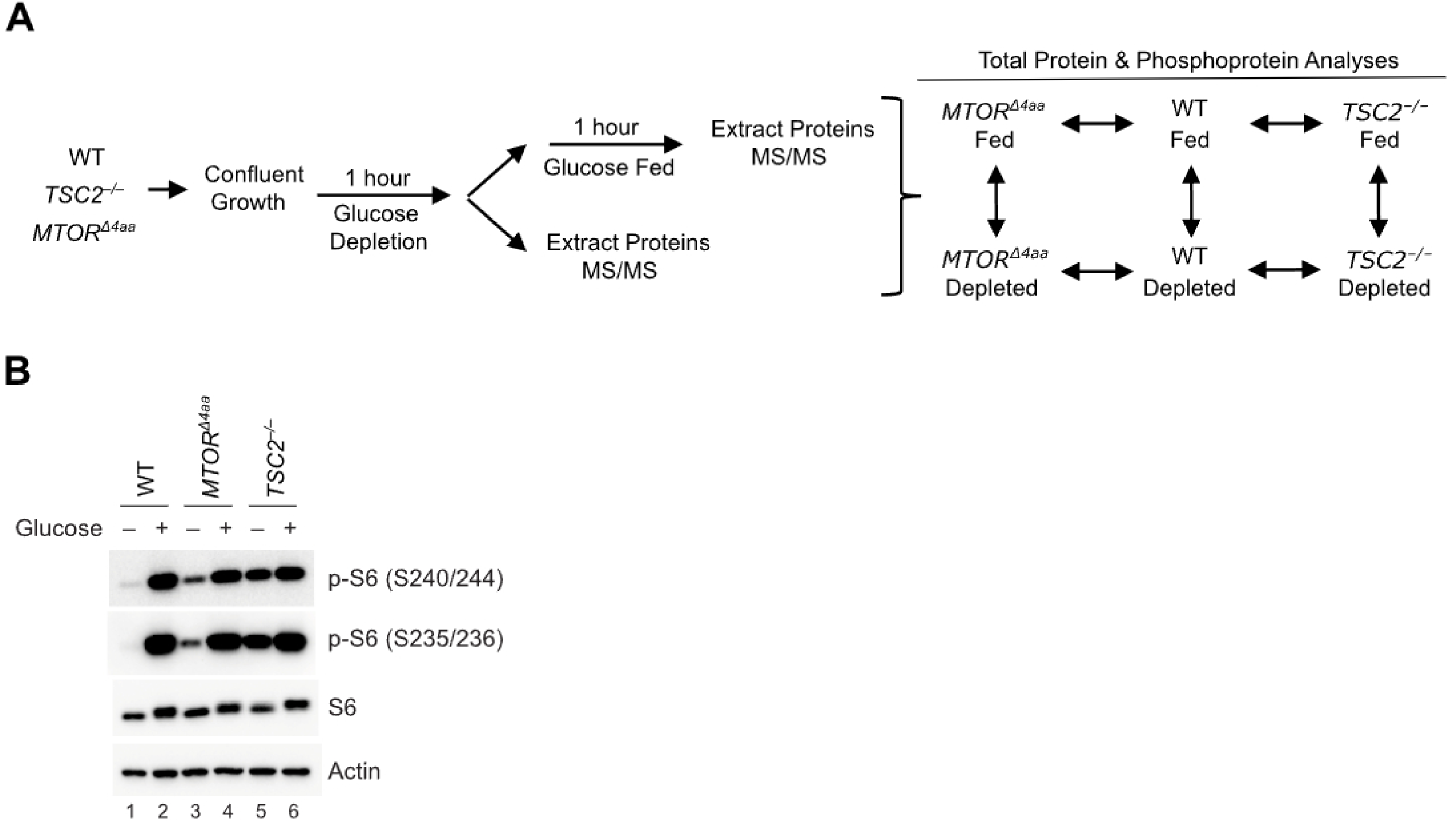
Distinct basal mTOR activity states in WT, *MTOR^Δ4aa^*, and *TSC2^−/−^* cells under glucose depletion. (A) Experimental scheme for proteomic and phosphoproteomic profiling (n=3). Arrows indicate pairwise comparisons for bioinformatic analysis. (B) WT, *MTOR^Δ4aa^* (R1480–C1483 deletion), and *TSC2^−/−^* U2OS cells were subjected to 1 h glucose depletion (−Glucose) followed by 1 h restimulation (+Glucose). Representative immunoblots show phospho-S6 (S240/244, S235/236), total S6, and actin loading controls. Basal mTOR activity under glucose depletion followed a graded hierarchy (*TSC2^−/−^* > *MTOR^Δ4aa^* > WT); restimulation induced mTOR activation in all genotypes.

Across all conditions, we identified over 6,000 proteins and 9,000 phosphosites with localization scores ≥0.75 across all samples (Figure S2A-D, Tables S1, S2). Biological triplicates showed tight clustering and high Pearson correlations (Figure S2E-H, S3A-B), confirming data quality and reproducibility. These comprehensive datasets provide the foundation for defining how the SKS mutation *MTOR^Δ4aa^* remodels the proteome and phosphoproteome across nutrient states and for distinguishing SKS-specific signaling signatures from classical mTORC1 hyperactivation.

### Distinct proteomic profiles in the glucose-depleted basal state distinguish MTOR^Δ4aa^ from TSC2^−/–^ cells

Under glucose depletion, mTORC1 activity is normally attenuated, allowing cells to conserve energy by reducing biosynthesis and increasing catabolic processes^39^. Because this nutrient-deprived condition places WT cells in a low-energy basal state, it provides an informative setting to distinguish whether SKS-associated mTOR activation resembles classical mTORC1 hyperactivation observed in TSC1/2 deficiency or instead adopts a distinct signaling profile. Having established that both *TSC2*^−/–^ and *MTOR^Δ4aa^* cells maintain elevated basal mTOR activity relative to WT based on the canonical target p-S6 (Figure 1B), we next asked how their proteomes compare under glucose depletion. We reasoned that if SKS mutations alter mTOR nutrient sensing through mechanisms distinct from loss of upstream TSC2-mediated inhibition, then *MTOR^Δ4aa^* cells should diverge not only from WT but also from *TSC2*^−/–^ cells in the basal state. Compared with WT, both *MTOR^Δ4aa^* and *TSC2*^−/–^ cells exhibited broad proteomic differences under glucose depletion (BH-adjusted p < 0.05; Figure 2A-B). *MTOR^Δ4aa^* cells displayed 1,014 upregulated and 964 downregulated proteins, while *TSC2^−/–^* cells showed an even greater magnitude of change (1,536 upregulated and 1,547 downregulated proteins), with some proteins reaching ∼5–6-fold change. Thus, under this glucose-depleted baseline condition, both mutant genotypes exhibited extensive proteomic remodeling relative to WT, consistent with persistent mTOR-linked signaling^39^.

**Figure 2.**
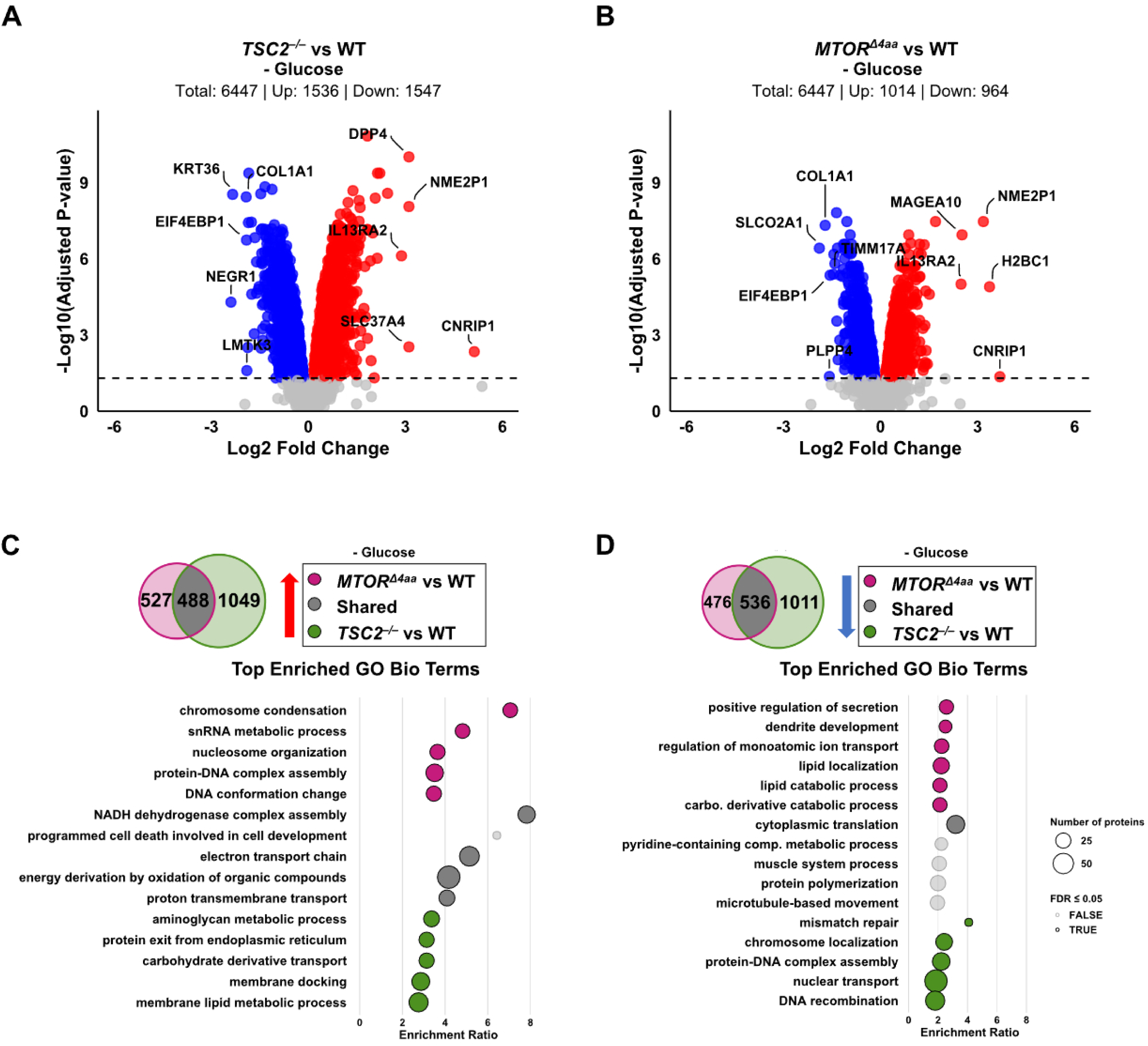
Shared basal metabolic activation but divergent proteomic programs in glucose-depleted *MTOR^Δ4aa^*and *TSC2^−/–^* cells. (A-B) Volcano plots illustrating protein expression changes under glucose depletion: (A) *MTOR^Δ4aa^* vs. WT cells and (B) *TSC2*^−/–^ vs. WT cells. Significantly up- (red) and down-regulated (blue) proteins (log_2_ fold change > 0, adjusted P-value ≤ 0.05) are shown; non-significant, gray. Largest fold-change proteins are labeled by gene name. (C-D) Venn diagrams (top) and GO Biological Process enrichment analysis (bottom) comparing differentially expressed proteins in *MTOR^Δ4aa^*against WT and *TSC2*^−/–^ against WT under glucose-depleted conditions. (C) Upregulated Proteins. (D) Downregulated proteins. Venn diagrams show overlap among significantly differentially expressed proteins (adjusted P ≤ 0.05). Bubble plots display enriched GO Biological Process terms (top 5 per category: *MTOR^Δ4aa^*unique, shared, and *TSC2*^−/–^ unique). Bubbles size presents the number of proteins mapped to each term and is shaded according to the False Discovery Rate (FDR) (solid bubbles: FDR ≤ 0.05; transparent bubbles: FDR > 0.05).

Overlap analyses revealed that 488 proteins were upregulated in both mutants relative to WT under glucose depletion and these proteins were primarily enriched for mitochondrial oxidative phosphorylation and NADH dehydrogenase complex assembly (Figure 2C, Table S3). This shared signature suggests persistent respiratory activity despite glucose deprivation, consistent with mTORC1-associated mitochondrial output^40,41^. Beyond this common metabolic core, each genotype displayed distinct remodeling patterns. Notably, *MTOR^Δ4aa^*cells uniquely upregulated nuclear processes associated with chromatin regulation and RNA processing, including chromosome condensation, nucleosome organization, snRNA metabolic processes, and protein-DNA complex assembly, suggesting persistent nuclear transcriptional and splicing activity^42^. In contrast, *TSC2*^−/–^ cells preferentially upregulated pathways related to membrane lipid metabolism, ER transport, and carbohydrate derivative metabolism, consistent with prior findings about its role in lipid synthesis and ER stress response^43,44^.

Downregulated proteins showed a similarly divided pattern (Figure 2D). A shared core of 536 downregulated proteins across both genotypes was enriched for cytoplasmic translation and protein polymerization, suggesting general translational restraint despite ongoing metabolic activity. However, *MTOR^Δ4aa^*cells selectively downregulated proteins involved in secretion, ion transport, and lipid catabolism. In contrast, *TSC2*^−/–^ cells downregulated nuclear transport, protein-DNA complex assembly, mismatch repair, and chromosomal localization factors. Notably, several of these downregulated pathways in *TSC2*^−/–^ cells were conversely upregulated in *MTOR^Δ4aa^*, underscoring divergent regulatory priorities of the two mutants.

Pathway-level enrichment analysis reinforced these distinctions. Both mutants upregulated electron transport chain and oxidative phosphorylation components (Figure S4A-B, Table S4). *MTOR^Δ4aa^*cells showed additional enrichment of DNA replication, RNA processing, and MAPK/ERK signaling, while downregulating the Parkin-mediated ubiquitination pathway and ferroptosis, suggesting altered mitochondrial quality control^45^. By contrast, *TSC2*^−/–^ cells uniquely upregulated sphingolipid metabolism and showed suppression of DNA repair and ribosomal protein pathways. Complex-level enrichment using the CORUM protein-complex database^24,25^ confirmed that both mutants upregulated mitochondrial complex I and 55S mitoribosome, whereas *MTOR^Δ4aa^* uniquely upregulated spliceosome complexes A and B under glucose depletion (Figure S5A-B, Table S5).

Together, these findings show that *MTOR^Δ4aa^*cells do not adopt a typical glucose depletion-adapted proteomic profile. Instead, they adopt a distinct program marked by persistent oxidative phosphorylation, chromatin regulation, and RNA-processing activity. This profile diverges from *TSC2^−/–^*cells, which also remain highly active under glucose depletion but are biased toward a more classically anabolic and metabolically imbalanced state. Thus, although both mutants deviate from a wild-type glucose-depletion response, *MTOR^Δ4aa^* cells exhibit a distinct baseline signaling architecture rather than phenocopying *TSC2* loss.

### Glucose restoration reinforces genotype-specific metabolic and gene-regulatory responses

Having established that *MTOR^Δ4aa^* and *TSC2*^−/–^ cells occupy distinct proteomic profiles under glucose depletion, we next asked how these mutants respond when glucose is restored. In WT cells, glucose refeeding normally re-engages mTORC1-dependent anabolic processes. We therefore reasoned that glucose restoration would reveal whether *MTOR^Δ4aa^* cells undergo broad anabolic remodeling similar to *TSC2*^−/–^ cells or instead reinforce the distinct gene-regulatory and metabolic programs already apparent under glucose depletion.

Compared with WT, both *MTOR^Δ4aa^* and *TSC2*^−/–^ cells exhibited broad proteomic differences following glucose refeeding (BH-adjusted p < 0.05; Figure 3A-B). *MTOR^Δ4aa^*cells showed 1,234 upregulated and 1,167 downregulated proteins relative to WT, while *TSC2*^−/–^ cells exhibited a comparable magnitude of change (1,080 upregulated and 1,048 downregulated proteins). Thus, both genotypes responded to glucose restoration but remained strongly divergent from WT, indicating that nutrient repletion does not normalize either mutant state.

**Figure 3.**
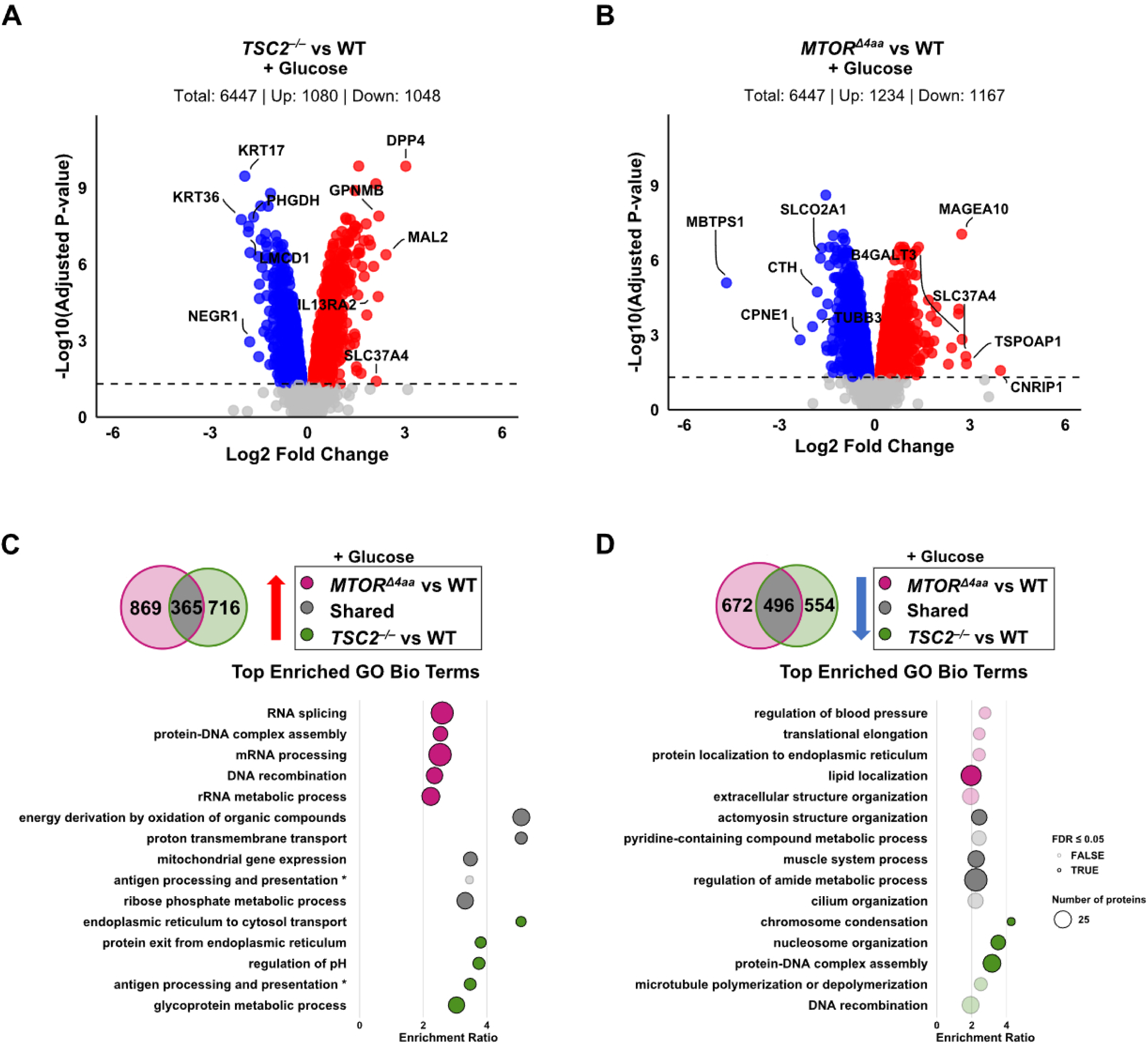
Glucose refeeding reinforces nuclear and transcriptional programs in *MTOR^Δ4aa^* cells but with limited anabolic remodeling. (A-B) Volcano plots illustrating protein expression changes. (A) Comparison of glucose-fed *MTOR^Δ4aa^*cells against glucose-fed wild-type (WT) cells. (B) Glucose-fed *TSC2*^−/–^ cells compared against glucose-fed WT cells. Color and labeling conventions as in Figure 2. (C-D) Venn diagrams (top) and GO Biological Process enrichment plots (bottom) comparing differentially expressed proteins in *MTOR^Δ4aa^* against (WT) and *TSC2*^−/–^ against WT under glucose-fed conditions. (C) Proteins upregulated in *MTOR^Δ4aa^* glucose-fed against WT glucose-fed and *TSC2*^−/–^ glucose-fed against WT glucose-fed cells. (D) Proteins downregulated in *MTOR^Δ4aa^* glucose-fed against WT glucose-fed and *TSC2*^−/–^ glucose-fed vs WT glucose-fed cells. Venn diagrams represent the overlap of significantly differentially expressed proteins (Adjusted P-Value ≤ 0.05; |Log_2_Fold Change| ≥ 0). Display conventions as in Figure 2.

Overlap analysis identified 365 proteins upregulated in both *MTOR^Δ4aa^*and *TSC2*^−/–^ cells relative to WT, while each genotype retained distinct sets of uniquely upregulated proteins (Figure 3C, Table S3). The shared proteins were strongly enriched for mitochondrial pathways, consistent with persistent mTOR-associated metabolic activation^40^. *MTOR^Δ4aa^* uniquely upregulated pathways related to RNA splicing, mRNA processing, and DNA recombination, whereas *TSC2*^−/–^ cells preferentially upregulated processes including ER transport, pH regulation, and glycoprotein processing.

Downregulated proteins showed a similarly divided pattern (Figure 3D). A shared set of 496 proteins was downregulated in both *MTOR^Δ4aa^*and *TSC2*^−/–^ cells compared with WT and was enriched for cytoskeletal and amide metabolic processes. *MTOR^Δ4aa^* cells uniquely downregulated pathways involved in lipid localization, whereas *TSC2*^−/–^ cells selectively downregulated chromatin-related processes, including DNA–rotein complex assembly and DNA recombination, which were conversely upregulated in *MTOR^Δ4aa^*. These reciprocal patterns further highlight that the two mutant genotypes engage distinct regulatory priorities even under nutrient-replete conditions.

Pathway-level enrichment analyses reinforced these genotype-specific differences. Both *MTOR^Δ4aa^* and *TSC2*^−/–^ cells upregulated the mitochondrial electron transport chain, consistent with shared activation of oxidative phosphorylation (Figure S4C-D, Table S4). However, *MTOR^Δ4aa^* cells additionally showed increased expression of mRNA processing, DNA replication, and DNA repair pathways. By contrast, *TSC2*^−/–^ cells displayed enrichment for sphingolipid metabolism and mitochondrial assembly pathways, consistent with reports linking TSC deficiency to altered sphingolipid regulation^43^. Further analysis using the CORUM protein complex database^24,25^ supported these patterns (Figure S5C-D, Table S5). *MTOR^Δ4aa^*cells upregulated the mitochondrial respiratory chain complex I, the mitochondrial 55S ribosome, and spliceosome complexes A, B, C, E, and 17S U2 snRNP^46^. *TSC2*^−/–^ cells also upregulated the mitochondrial respiratory chain complex I and the mitochondrial 55S ribosome, with additional enrichment of COX1 assembly factors. Both genotypes showed downregulation of the Parkin ubiquitination degradation pathway, suggesting shared alterations in mitochondrial quality control^47^.

Together, these findings indicate that glucose restoration reinforces, rather than erases, the distinct proteomic profiles established under glucose depletion. Across both depletion and refeeding conditions, *MTOR^Δ4aa^* and *TSC2*^−/–^ cells maintain elevated mitochondrial oxidative phosphorylation and reduced amide metabolism and cytoplasmic translation, consistent with sustained energy production coupled with restrained bulk protein synthesis. Beyond this shared metabolic core, the two mutants diverge markedly. *MTOR^Δ4aa^*cells preferentially engage nuclear programs, including protein–DNA complex assembly, RNA processing, and chromatin-associated remodeling, with reduced ER and lipid pathway activity. In contrast, *TSC2*^−/–^ cells favor ER-, membrane-, and lipid-associated processes, with reduced nuclear regulatory activity. Thus, despite similar mitochondrial activation, the two mutants allocate cellular resources in opposite directions: nuclear regulation in *MTOR^Δ4aa^* cells vs ER–lipid biosynthesis in *TSC2*^−/–^ cells.

### MTOR^Δ4aa^ cells exhibit a constrained dynamic range across glucose states

Next, we compared glucose-depleted and glucose-fed states within each genotype to evaluate their proteomic dynamic range following nutrient stimulation. Our data show that WT cells displayed a robust glucose-responsive shift, with 253 upregulated and 204 downregulated proteins (BH-adjusted p < 0.05; Figure 4A, Table S1). *TSC2*^−/–^ cells showed an even broader response to glucose, with 392 upregulated and 322 downregulated proteins (BH-adjusted p < 0.05; Figure 4B), indicating exaggerated nutrient responsiveness despite constitutive mTOR signaling^39^. Strikingly, *MTOR^Δ4aa^* cells resembled neither WT nor *TSC2*^−/–^. These cells showed only 63 upregulated and 35 downregulated proteins (Figure 4C), indicating a constrained proteomic dynamic range or lack of metabolic plasticity in response to glucose restimulation. Despite these quantitative differences, a small core group of nutrient-responsive proteins was shared across genotypes, with 15 upregulated and 7 downregulated proteins (Figure 4A-C). The shared upregulated proteins included canonical immediate-early transcription factors (FOS, EGR1, NR4A1), as well as CCN2 (aka CTGF), that are rapidly induced by growth stimuli^48^, indicating that early stimulus sensing remains intact. Shared downregulated proteins included cytoskeletal regulators PDLIM5 and TNS1, as well as the receptor tyrosine kinase AXL.

**Figure 4.**
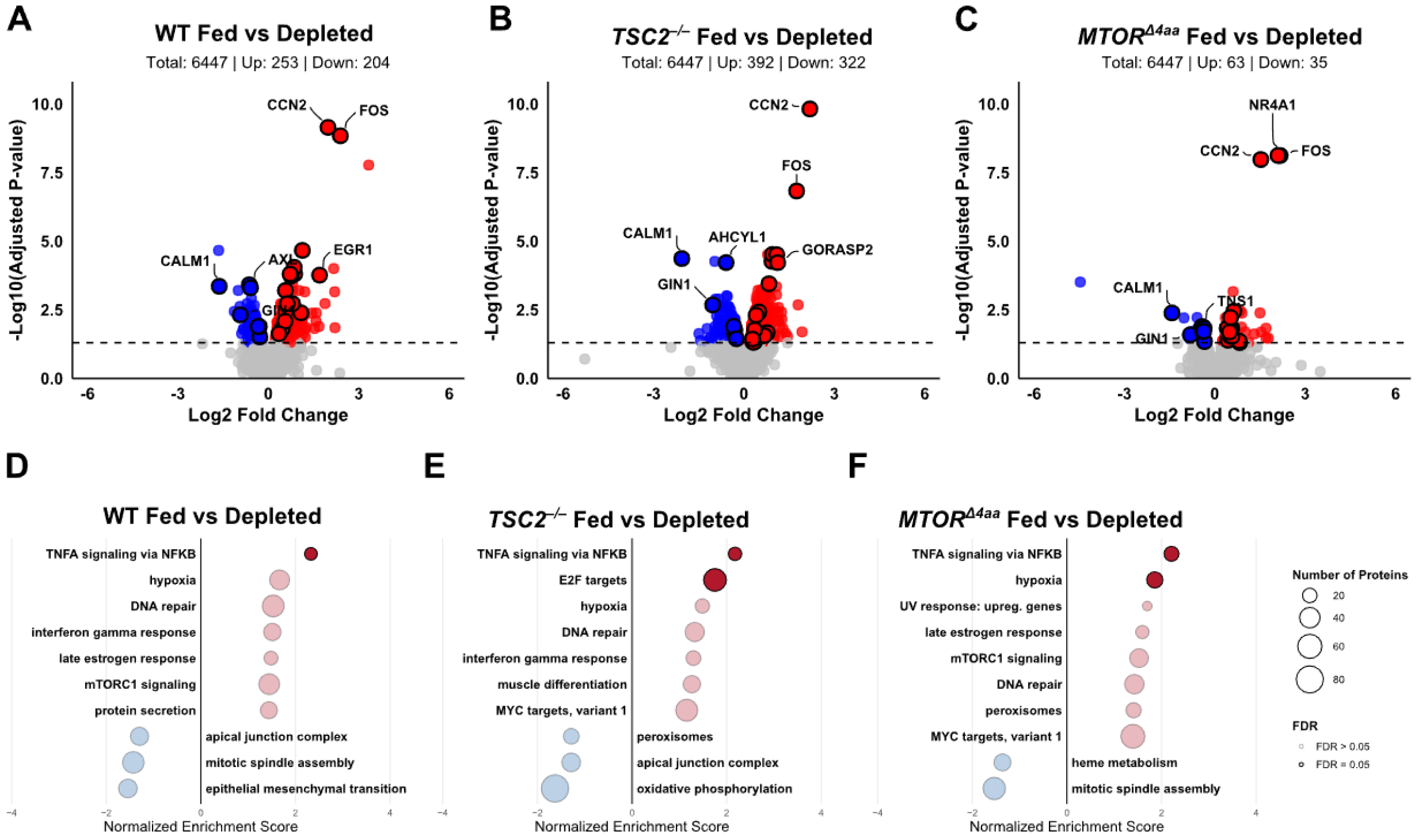
*MTOR^Δ4aa^* cells exhibit constrained proteome remodeling upon glucose stimulation. (A-C) Volcano plots illustrating protein expression changes. (A) Comparison of glucose-fed WT cells against glucose-depleted WT cells. (B) Glucose-fed *TSC2*^−/–^ cells compared against glucose-depleted *TSC2*^−/–^ cells. (C) Glucose-fed *MTOR^Δ4aa^* cells compared against glucose-depleted *MTOR^Δ4aa^*cells. Color conventions as in Figure 2. Significantly regulated proteins shared across all three genotypes are outlined in black and labels denote the top five shared up- and down-regulated proteins in each comparison, ranked by Log_2_ Fold Change with gene names. (D-F) Corresponding Gene Set Enrichment Analysis (GSEA) results using the MSigDB Hallmark Gene Set database for (D) glucose-fed WT cells compared against glucose-depleted WT cells, (E) glucose-fed *TSC2*^−/–^ cells compared against glucose-depleted *TSC2*^−/–^ cells, and (F) glucose-fed *MTOR^Δ4aa^* cells compared against glucose-depleted *MTOR^Δ4aa^* cells. Each plot displays the top 10 enriched gene sets ranked by normalized enrichment score derived from Log_2_ Fold Change values. Bubbles are sized by the number of proteins mapped to each gene set, colored by direction of enrichment (red = upregulated, blue = downregulated), and shaded by False Discovery Rate (solid bubbles = FDR ≤ 0.05; transparent bubbles = FDR > 0.05).

To identify pathways underlying these genotype-specific proteomic shifts, we performed gene set enrichment analysis (GSEA) using MSigDB hallmark gene sets^22^. All three genotypes showed comparable positive enrichment for TNFα signaling via NF-κB upon glucose refeeding, with the enrichment driven largely by immediate early genes (FOS, EGR1, NR4A1) characteristic of a generic nutrient-responsive transcriptional program (Figure 4D-F, Table S6). Interferon gamma response was additionally enriched in WT and *TSC2*^−/–^ cells (FDR < 0.05; Figure 4D-F). Beyond this shared early response signature, the broader responses diverged substantially across genotypes. WT cells mounted a coordinated nutrient response, with enrichment of mTORC1 signaling, translational and secretory pathways, and hormone-responsive transcriptional programs, consistent with reactivation of anabolic and biosynthetic processes upon glucose restoration (Figure 4D). In contrast, *TSC2*^−/–^ cells showed enrichment of E2F and MYC targets, accompanied by suppression of oxidative metabolism (Figure 4E). This pattern suggests a growth-biased transcriptional response, consistent with constitutive mTOR activation driving cell-cycle-associated programs rather than balanced metabolic remodeling.

By comparison, *MTOR^Δ4aa^* cells retained the immediate early and hypoxia-associated response signatures but lacked broader anabolic or metabolic remodeling in response to glucose restimulation (Figure 4F). In contrast to WT and *TSC2*^−/–^ cells, pathways linked to translation, secretion, and metabolic remodeling were not strongly induced in *MTOR^Δ4aa^* cells. Notably, peroxisomal metabolism trended upward in *MTOR^Δ4aa^* but was repressed in *TSC2*^−/–^, suggesting that *MTOR^Δ4aa^* cells retain aspects of redox- and lipid-associated programs while remaining comparatively restrained in response to glucose cues.

Together, these findings indicate that WT cells execute a coordinated metabolic switch upon glucose restoration, whereas *TSC2*^−/–^ cells exhibit a growth-biased, dysregulated response, and *MTOR^Δ4aa^* cells instead occupy a constrained state within a narrower proteomic range than WT or *TSC2*^−/–^ cells. Combined with the distinct depletion and refeeding profiles, this reduced dynamic range suggests that *MTOR^Δ4aa^*cells occupy a partially desensitized signaling state in which proximal nutrient-responsive programs are retained, but the capacity for downstream anabolic and metabolic remodeling in response to nutrient and energy cues is limited.

### Transcriptional profiling reveals distinct but overlapping basal regulatory states in MTOR^Δ4aa^ and TSC2^−/–^cells

mTOR signaling has been implicated in coordinating transcriptional responses to metabolic stress^49–51^. To characterize the transcriptional programs underlying the basal proteomic differences observed between genotypes, we performed RNA-seq under glucose depletion conditions (Figure S6A, lanes 7-9 selected for RNA-seq). Each pairwise comparison identified ∼3,000-4,000 differentially expressed genes per direction (Table S7; BH-adjusted p < 0.05), confirming that each genotype occupies a transcriptionally distinct state. Transcription factor (TF) activity inference revealed strong enrichment of inflammatory-associated TFs in *TSC2*^−/–^ cells, including REL, NFKB1, RELA, STAT2, IRF1, IRF9, and RELB, alongside suppression of E2F4, TFDP1, and SMAD5 (Figure S6B, Table S8). *MTOR^Δ4aa^* cells shared enrichment of REL, RELA, NFKB1, STAT2, and IRF1, though at lower magnitude, but otherwise diverged, showing strongest activation of RFX5, ATF3, and MYC, with suppression of TEAD1, MEF2A, RUNX2, and TCF12 (Figure S6C, Table S8). Direct comparisons confirmed this inversion: MYC, E2F family members, ATF3, and MYB were enriched in *MTOR^Δ4aa^* cells, while STAT1, ESRRA, HIF1A, and SREBF1/2 were enriched in *TSC2*^−/–^ (Figure S6D, Table S8). GSEA was consistent with these profiles: *TSC2*^−/–^ cells were enriched for interferon gamma response, while *MTOR^Δ4aa^* cells were enriched for E2F targets, MYC targets, and G2/M programs, mirroring their respective proteomic biases toward ER/lipid and nuclear/RNA-processing programs, respectively (Figure S6E-G, Table S9). Notably, canonical mTORC1 transcriptional signatures were reduced in *MTOR^Δ4aa^*cells despite elevated kinase activity, suggesting partial uncoupling of mTOR activation from downstream transcriptional outputs.

### Distinct nutrient-state kinase signatures define divergent signaling in MTOR^Δ4aa^ cells

mTOR signaling rapidly remodels kinase networks as cells transition across nutrient states. Under glucose depletion, energy-sensing kinases such as AMPK suppress mTORC1 activity to conserve energy^37^, whereas glucose restoration re-engages growth-associated pathways, including mTOR and ERK/MAPK^39,48^. Because kinase signaling networks sit upstream of transcription control, these transitions are major determinants of the transcriptional programs during nutrient stress and recovery. Having established that *MTOR^Δ4aa^* and *TSC2*^−/–^ cells occupy distinct transcriptional and proteomic states, we next asked whether these differences are reflected in their underlying phospho-signaling profiles. Because *TSC2* loss removes upstream inhibition of mTORC1, whereas SKS mutations alter mTOR itself, we reasoned that these two hyperactive mTOR states would engage distinct kinase networks across nutrient conditions.

Under glucose depletion, both *TSC2*^−/–^ and *MTOR^Δ4aa^*cells displayed extensive phosphoproteomic differences relative to WT cells (182 up/152 down and 158 up/118 down, respectively; BH-adjusted p < 0.05; Figure 5A-B, Table S2). Despite similar magnitudes of change, kinase enrichment analysis derived from substrate-motif inference revealed different signaling profiles (Figure 5C-D, Table S10). *TSC2*^−/–^ cells maintained activation of AGC-family kinases (AKT2, ROCK1, SGK3) that are known to mediate mTOR-driven anabolic growth, survival, and metabolism, along with suppression of AMPKA2 and LKB1. This is notable given recent evidence that mTORC1 can directly inhibit AMPK signaling even under nutrient stress conditions^52^. In contrast, *MTOR^Δ4aa^* cells showed broad suppression across major kinase families, including AGC kinases involved in translational and metabolic control (PDPK1, MASTL), CAMKs linked to stress- and calcium-responsive signaling (MAPKAPK2/3/5, BRSK1/2, MELK), and STE/TKL kinases that drive MAPK cascades (TAK1, MLK2, MEKK1, HPK1, MST1/3). Activity was retained only in a limited subset of AGC and CAMK kinases associated with cytoskeletal and contractile regulation (MRCKA, CRIK, CAMLCK) (Figure 5C). Among tyrosine kinases, *MTOR^Δ4aa^* cells showed selective activation of JAK2 and MST1R, with suppression of JAK1 and DDR1 (Figure 5D, Table S11). Notably, canonical stress- and inflammation-associated kinases, including IRAK1, TAK1, and MAPKAPK2/3/5, were suppressed rather than activated under basal conditions in *MTOR^Δ4aa^*cells. The LKB1-AMPK axis, a central energy stress pathway, was also suppressed in *MTOR^Δ4aa^* cells, indicating impaired engagement of canonical energy-sensing responses. Kinase activity profiles were largely non-overlapping between *MTOR^Δ4aa^* and *TSC2*^−/–^ cells under glucose depletion relative to WT, underscoring different signaling strategies: broader constitutive kinase engagement in *TSC2*^−/–^ versus more selective and restrained signaling in *MTOR^Δ4aa^*.

**Figure 5.**
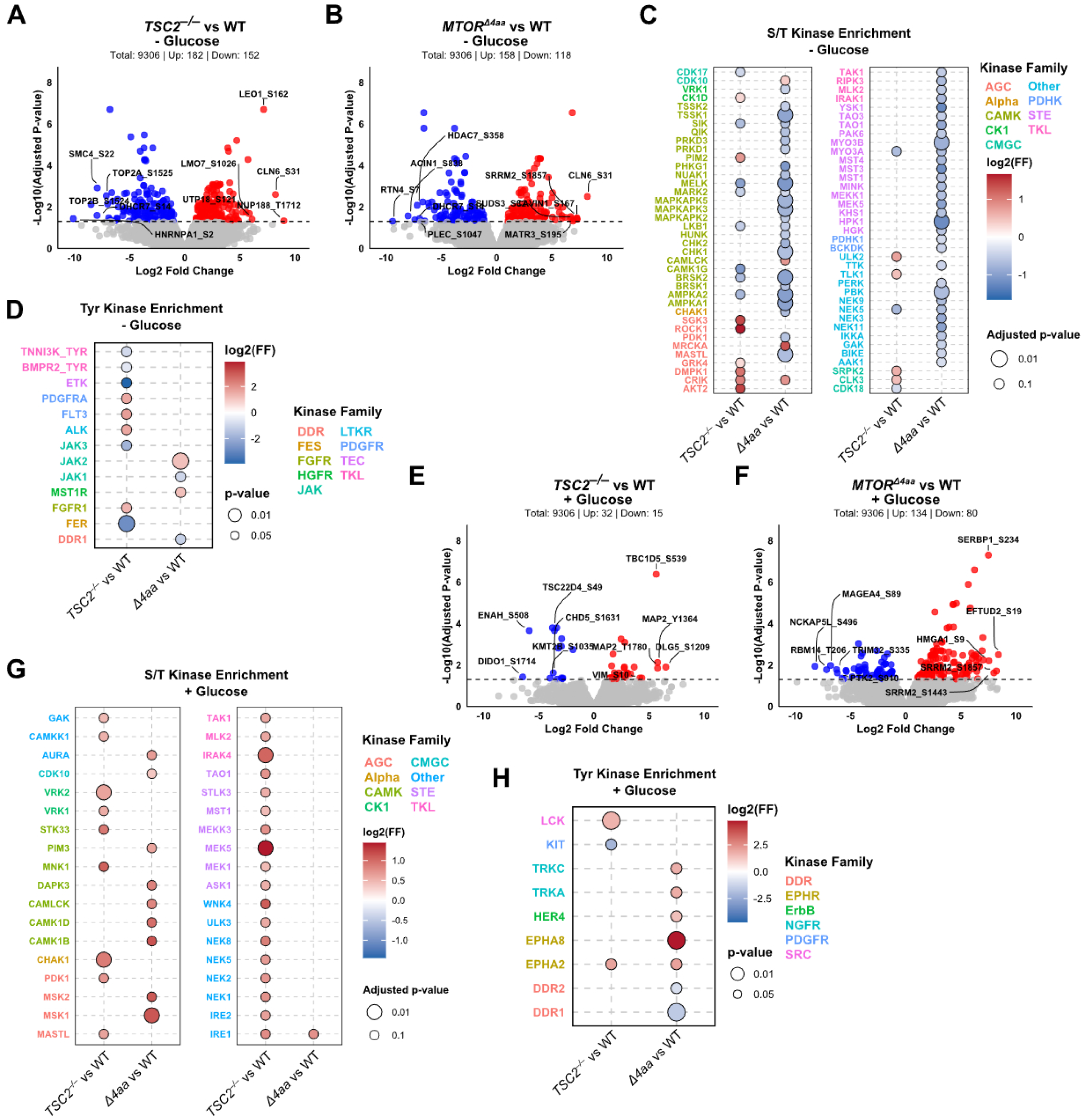
Distinct nutrient-state phosphoproteomic profiles in *MTOR^Δ4aa^* and *TSC2 ^-/-^* cells. (A-B) Volcano plots showing changes in phosphosite changes under glucose depletion: (A) *TSC2*^−/–^ vs WT and (B) *MTOR^Δ4aa^* vs WT. Significantly upregulated phosphosites (Log_2_ Fold Change > 0; Adjusted P-value ≤ 0.05) are shown in red; downregulated (Log_2_ Fold Change < 0; Adjusted P-value ≤ 0.05) in blue; nonsignificant in gray. Top phosphosites are labeled by gene and residue. (C-D) Kinase enrichment analysis under glucose depletion. (C) Serine/threonine kinases; (D) tyrosine kinases. Comparisons include *TSC2*^−/–^ vs WT and *MTOR^Δ4aa^*vs WT. The x-axis shows genotype comparisons; the y-axis lists kinases grouped by family. Bubbles color represents z-scored enrichment (blue = activated, red = inhibited); size reflects adjusted P-value. (E-F) Volcano plots showing phosphosite changes under glucose refeeding: (E) *TSC2*^−/–^ vs WT and (F) *MTOR^Δ4aa^* vs WT. (G-H) Kinase enrichment under glucose refeeding. (G) Serine/threonine kinases; (H) tyrosine kinases. Display conventions same as in (C-D).

Upon glucose refeeding, these differences became even more pronounced. *MTOR^Δ4aa^* cells exhibited a substantial phosphoproteomic shift (134 up/80 down; Figure 5E, Table S10), whereas *TSC2*^−/–^ cells showed a comparatively modest response (32 up/15 down; Figure 5F). Kinase enrichment revealed that *MTOR^Δ4aa^*cells preferentially engaged MAPK/ERK-linked kinases (MSK1/2; downstream effectors linking MAPK signaling to transcription), Ca^2+^/CaM-regulated kinases (CAMK1B/D, CAMLCK, DAPK3; activity-dependent signaling pathways), as well as PIM3 and AURA, which are associated with transcriptional regulation and cytoskeletal remodeling (Figure 5G). MSK1/2 activation may provide a mechanistic link between the ERK/CaMK kinase signatures and E2F/MYC-driven transcriptional identity observed in *MTOR^Δ4aa^* cells. This pattern indicates that nutrient-responsive signaling in *MTOR^Δ4aa^* cells is preferentially routed toward ERK and Ca^2+^/CaM-dependent pathways, pathways that couple environmental and activity signals to gene regulation, rather than canonical mTORC1-mediated translational control. By contrast, *TSC2*^−/–^ cells re-engaged a broad set of growth-associated kinases, including PDK1 (a master activator of AGC kinases downstream of PI3K signaling) and MAPK pathway kinases (MEK1/5, TAK1), consistent with coordinated mTOR- and MAPK-associated anabolic and proliferative signaling. These differences were also reflected in tyrosine kinase activities: *MTOR^Δ4aa^* selectively activated EPHA2/EPHA8 and HER4 (receptor tyrosine kinases feeding into PI3K-AKT-mTOR and MAPK pathways) and suppressed DDR1/2 (matrix-associated signaling) relative to WT. *TSC2*^−/–^ cells showed a distinct pattern marked by activation of LCK and EPHA2 (Figure 5H, Table S11), further highlighting divergent extracellular signaling integration.

Together, these findings indicate that *MTOR^Δ4aa^* and *TSC2*^−/–^ cells engage distinct kinase networks at both basal and stimulated states. Under glucose depletion, *MTOR^Δ4aa^* cells exhibit impaired AMPK engagement and selectively retain stress- and adhesion-associated kinases. *TSC2*^−/–^ cells also suppress AMPK, yet maintain broader growth- and metabolism-linked kinase activity. Upon glucose restoration, these differences persist: *MTOR^Δ4aa^*preferentially activates MAPK- and CaMK-associated signaling pathways, while *TSC2*^−/–^ re-engages a growth-driven anabolic kinase network centered on AGC and MAPK pathways. These results suggest that *MTOR^Δ4aa^*cells occupy a rewired signaling state, rather than a simple amplification of canonical mTORC1 activity.

### Glucose refeeding amplifies MAPK and CaMK signaling in MTOR^Δ4aa^ cells

Consistent with the distinct kinase architectures observed under glucose depletion, we next asked how these signaling networks are reconfigured upon nutrient restoration. mTOR coordinates energy sensing and cellular metabolism through both transcriptional^49–51,53^ and translational control^54^. While proteomic changes reflect longer-term shifts in protein abundance, phosphorylation dynamics capture rapid signaling responses to nutrient shifts. Upon nutrient restoration, mTORC1 and mTORC2 typically re-engage anabolic growth via S6K1, AKT, and 4E-BP1, as well as transient MAPK/ERK signaling to support transcriptional response^55–57^. Given that *MTOR^Δ4aa^* cells maintain persistent nuclear and mitochondrial programs across glucose states relative to WT, we asked whether glucose refeeding elicits a distinct phospho-signaling response within each genotype.

In WT cells, glucose refeeding induced phosphoproteomic remodeling (132 up, 184 down; BH-adjusted p < 0.05; Figure 6A, Table S2). *TSC2*^−/–^ cells exhibited a comparable response (126 up, 113 down; Figure 6B). In contrast, *MTOR^Δ4aa^* cells showed the largest shift (198 up, 171 down) (Figure 6C), despite relatively limited proteomic changes between glucose states (Figure 4C). Thus, *MTOR^Δ4aa^* cells display disproportionate phosphorylation changes relative to changes in protein abundance.

**Figure 6.**
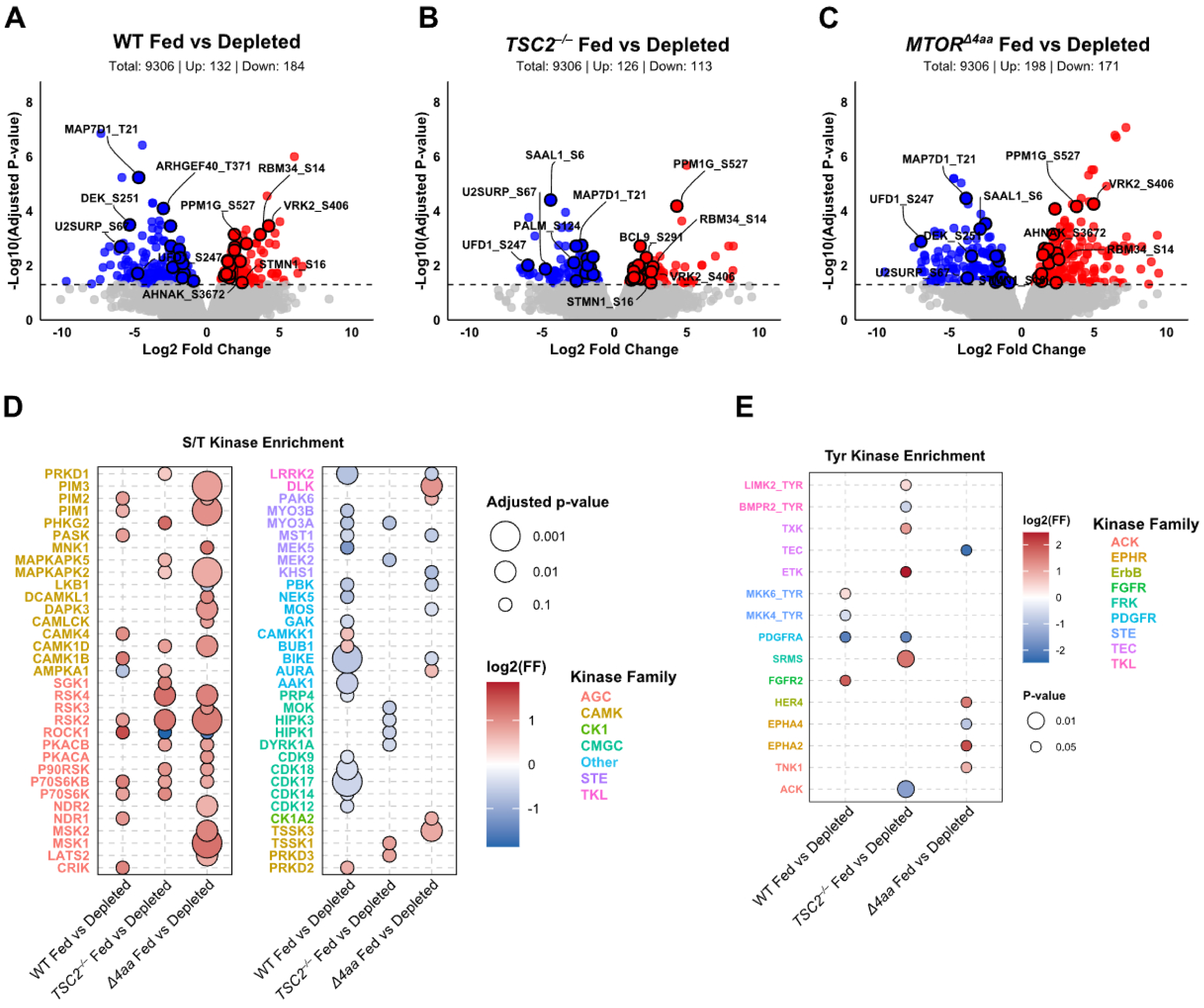
Glucose refeeding selectively amplifies MAPK/ERK and CaMK signaling in *MTOR^Δ4aa^* cells. (A-C) Volcano plots illustrating phosphosite expression changes. (A) Comparison of glucose-fed WT cells against glucose-depleted WT cells. (B) Glucose-fed *TSC2*^−/–^ cells compared against glucose-depleted *TSC2*^−/–^ cells. (C) Glucose-fed *MTOR^Δ4aa^* cells compared against glucose-depleted *MTOR^Δ4aa^* cells. Significantly upregulated phosphosites (Log_2_ Fold Change > 0; Adjusted P-value ≤ 0.05) are shown in red; significantly downregulated phosphosites (Log_2_ Fold Change < 0; Adjusted P-value ≤ 0.05) are shown in blue; nonsignificant phosphosites are shown in gray. Significantly regulated phosphosites shared across all three genotypes are outlined in black, and phosphosites with the largest fold change are labeled with gene names + phosphosite. (D) Serine/Threonine (S/T) kinase enrichment bubble plot comparing glucose-fed WT cells against glucose-depleted WT cells, glucose-fed *TSC2*^−/–^ cells against glucose-depleted *TSC2*^−/–^ cells, and glucose-fed *MTOR^Δ4aa^* cells against glucose-depleted *MTOR^Δ4aa^* cells, and. The x-axis shows the genotype and nutrient-status comparison; the y-axis lists kinases color-coded by kinase family. Bubbles are colored by z-scored Log_2_ Enrichment (FF) (blue = activated, red = inhibited), and sized by adjusted p-value. (E) Tyrosine (Tyr) kinase enrichment bubble plot comparing glucose-fed WT cells against glucose-depleted WT cells, glucose-fed *TSC2*^−/–^ cells against glucose-depleted *TSC2*^−/–^ cells, and glucose-fed *MTOR^Δ4aa^* cells against glucose-depleted *MTOR^Δ4aa^*cells. The x-axis shows the genotype and nutrient-status comparison; the y-axis lists kinases color-coded by kinase family. Bubbles are colored by z-scored Log_2_ Enrichment (FF) (blue = activated, red = inhibited), and sized by p-value.

Kinase enrichment analysis further distinguished the three genotypes (Figure 6D, Table S10). In WT cells, glucose feeding reactivated S6K1 and S6K2 and suppressed AMPK, consistent with restoration of anabolic signaling and relief of energy stress. *TSC2*^−/–^ cells likewise activated several AGC-family kinases, including S6K1, S6K2, RSK2/4, P90RSK, and SGK1. In contrast to WT cells, however, AMPK was not suppressed but rather induced upon refeeding, consistent with persistent energetic stress despite glucose availabillity^58^. In contrast, *MTOR^Δ4aa^* cells displayed a distinct kinase signature and preferentially engaged ERK effectors (RSK, MSK, and MNK1; link MAPK signaling to transcription and translation), MAPK-activated kinase MAPKAPK2 (stress and inflammatory signaling), Ca^2+^/CaM-dependent kinases (CAMK1D, CAMLCK; activity- and calcium-dependent signaling), and PIM kinases (PIM1/2/3), which provide an alternative route to sustain translational output through phosphorylation of 4E-BP1 and PRAS40^59^ (Figure 6D). Notably, S6K1 did not exhibit significant nutrient-responsive activation in *MTOR^Δ4aa^* cells (Figure 7D), indicating limited engagement of canonical mTORC1 signaling during glucose refeeding. Given that *MTOR^Δ4aa^* cells exhibit elevated basal p-S6 under glucose depletion (Figure 1B, S6A), this likely reflects reduced dynamic range rather than absence of mTOR signaling. Together, these observations suggest that *MTOR^Δ4aa^* cells preferentially engages MAPK/ERK- and Ca^2+^/CaMK- dependent pathways that couple environmental and intracellular signals to transcriptional regulation, while canonical S6K1-associated substrate enrichment remains limited during this nutrient state transition.

**Figure 7.**
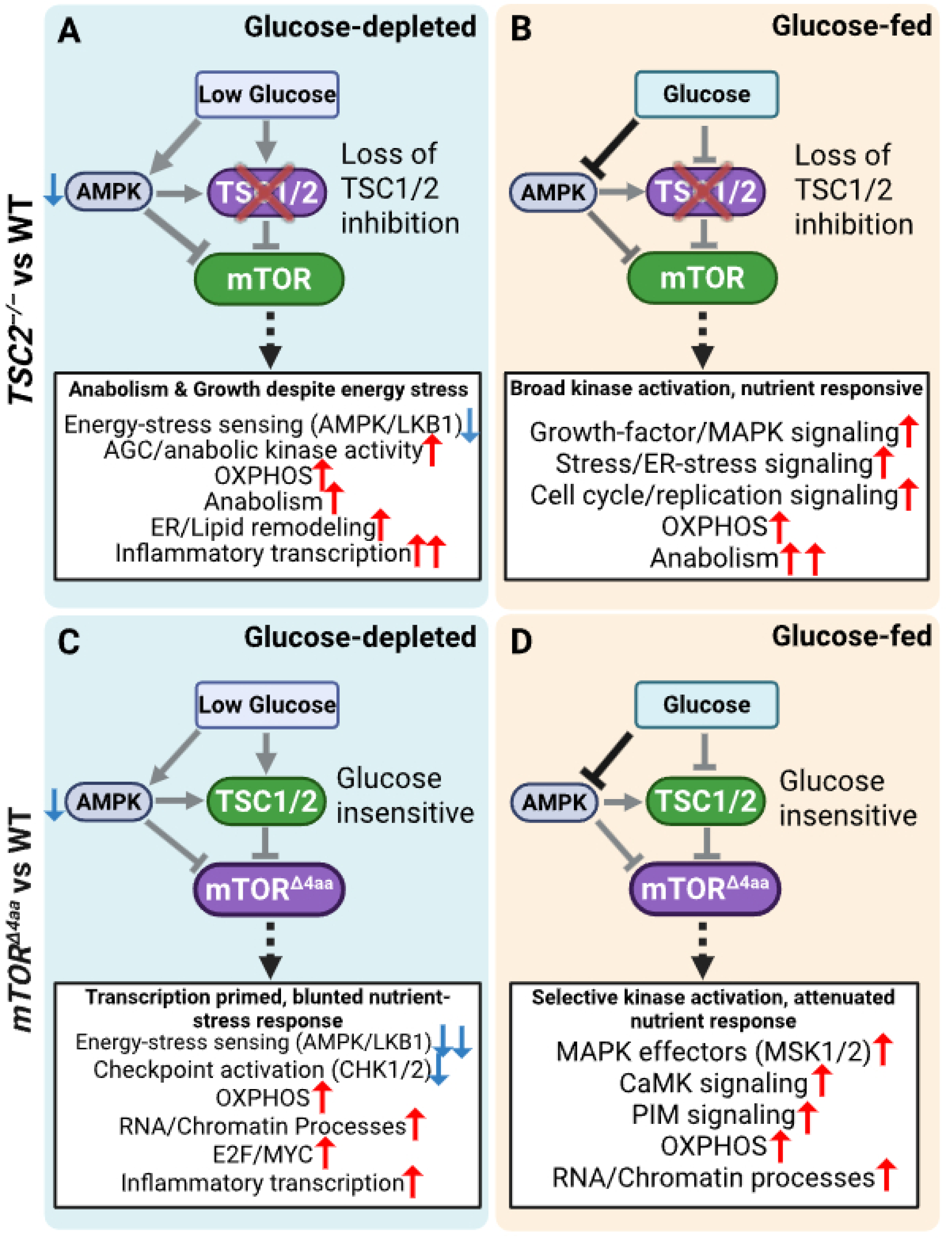
Model of distinct mTOR network states in *MTOR^Δ4aa^* and *TSC2^−/–^* cells. (A) Under glucose depletion, *TSC2*^−/–^ cells sustain anabolism despite energy stress, with elevated AGC/anabolic kinase activity, OXPHOS, ER/lipid remodeling, and inflammatory transcription, alongside attenuated energy-stress sensing (AMPK/LKB1). (B) Upon refeeding, *TSC2*^−/–^ cells engage a broad kinase network including growth factor/MAPK, stress, and cell-cycle/replication-associated signaling, with further elevation of OXPHOS and anabolic programs. (C) Under glucose depletion, *MTOR^Δ4aa^* cells display blunted energy-response signaling, failing to engage energy-stress sensing (AMPK/LKB1), checkpoint activation (CHK1/2), and ER stress signaling (PERK), while maintaining elevated OXPHOS, RNA/chromatin processes, E2F/MYC transcription, and inflammatory transcription. (D) Upon refeeding, *MTOR^Δ4aa^* cells show selective kinase activation, including MAPK effectors (MSK1/2), CaMK signaling, and PIM signaling, with sustained OXPHOS and RNA/chromatin processes but limited additional anabolic remodeling. Solid black arrows, active regulatory interaction; grey arrows, attenuated/uncoupled interactions relative to WT; dashed arrows, cellular states inferred from proteomic and kinase enrichment analyses (not direct mTOR phosphorylation). Red and blue indicate increases and decreases relative to WT, respectively.

Tyrosine kinase enrichment further highlighted these differences (Figure 6E, Table S11). In WT cells, FGFR2 and MKK6 (a dual-specificity kinase upstream of MAPK pathways) were selectively activated, consistent with MAPK activation during nutrient refeeding^60^. In *TSC2*^−/–^ cells, activation of non-receptor tyrosine kinases (SRMS, ETK, TXK) reflects broad intracellular signaling. In contrast, *MTOR^Δ4aa^* cells preferentially activated receptor tyrosine kinases EPHA2 and HER4, which feed into PI3K-AKT-mTOR and MAPK pathways^61,62^. These patterns further support a signaling architecture in *MTOR^Δ4aa^*cells that emphasizes upstream receptor and ERK-linked signaling inputs rather than a coordinated activation of canonical mTORC1 outputs.

Together, these results demonstrate that glucose refeeding produces a hyper-responsive yet selectively rewired phosphorylation network in *MTOR^Δ4aa^* cells. While ERK/MAPK- and Ca^2+^/CaM -linked kinases that couple signaling to transcription are prominently engaged, coordinated enrichment of canonical S6K1-associated signaling is limited. This signaling configuration is consistent with the transcriptional and proteomic data, in which *MTOR^Δ4aa^* cells prioritize nuclear and gene-regulatory programs, whereas *TSC2*^−/–^ cells engage broader anabolic remodeling. Across nutrient states, *MTOR^Δ4aa^* and *TSC2*^−/–^ cells occupy distinct kinase landscapes: *TSC2*^−/–^ cells exhibit broad and uniformly engaged canonical mTORC1-driven anabolic signaling, whereas *MTOR^Δ4aa^* cells preferentially engage ERK- and CaMK-driven gene expression programs (Figure 7). This phosphoproteomic divergence mirrors their proteomic profiles and supports the view that SKS represents a distinct mode of mTOR dysregulation that differs from TSC2 deficiency as in TSC.

## Discussion

### Isogenic cell models enable direct comparison of SKS and TSC signaling states

In this study, we examined how a SKS mutation alters mTOR signaling by comparing isogenic CRISPR-engineered U2OS cell models of SKS (*MTOR^Δ4aa^*), classical mTORC1 hyperactivation (*TSC2*^−/–^), and WT cells under glucose depletion and refeeding. This system enables direct comparison of distinct hyperactive mTOR states in a controlled genetic background, providing an important foundation for mechanistic understanding of SKS-associated signaling changes. Several SKS variants have been characterized as gain-of-function based on increased phosphorylation of canonical mTOR substrates^3–6^. However, these measurements capture only a limited subset of mTOR outputs. The present study was designed to examine whether SKS mutations phenocopy classical mTORC1 hyperactivation or instead rewire signaling more broadly across multi-omics and molecular layers.

### MTOR^Δ4aa^ adopts a nutrient-desynchronized signaling profile

Our working model summarizes the differences between *MTOR^Δ4aa^* and *TSC2*^−/–^ cells (Figure 7). We show that *MTOR^Δ4aa^* cells exhibit a nutrient-desynchronized signaling profile distinct from *TSC2*^−/–^ cells, supporting the view that mTOR functions as a modular signaling network rather than a linear cascade^63^. Although both mutants shared upregulated oxidative phosphorylation and mitochondrial respiration, *MTOR^Δ4aa^* cells display a markedly constrained proteomic dynamic range across glucose states, in contrast to the broader remodeling observed *TSC2*^−/–^ cells. At the signaling level, *MTOR^Δ4aa^* cells preferentially engage MAPK/ERK, CAMK, and PIM pathways, while canonical mTORC1/S6K1 displays limited nutrient responsiveness. In parallel, *MTOR^Δ4aa^*cells maintain enriched nuclear and gene-regulatory programs, including RNA processing, chromatin regulation, and DNA repair, whereas *TSC2*^−/–^ cells are biased toward ER-, lipid-, and metabolic remodeling. Transcriptomic profiling further refines this distinction: both mutants share a baseline of cytokine-associated signaling, but diverge in downstream regulatory programs. *TSC2*^−/–^ cells are more strongly organized around NF-κB-, STAT-, and interferon-responsive transcription, whereas *MTOR^Δ4aa^* cells are enriched for E2F- and MYC-driven programs despite reduced canonical mTORC1 transcriptional signatures. Together, these findings indicate that the *MTOR^Δ4aa^*mutation does not simply increase mTORC1 activity but instead compresses nutrient-responsive signaling and redistributes downstream outputs toward transcriptional and chromatin-associated regulation. In contrast, *TSC2* deficiency sustains broader anabolic remodeling and canonical AGC-family kinase engagement.

### SKS exhibits altered mTOR coupling to nutrient sensing and RNA regulation

*TSC2* loss removes upstream inhibition of RHEB, enabling constitutive activation of mTORC1 and robust phosphorylation of canonical targets such as S6K1 and 4E-BP1 largely independent of nutrient status^33^. In contrast, SKS mutations occur within the mTOR kinase itself, clustering in the FAT and kinase domains^4^. The FAT domain coordinates conformational changes required for RHEB-mediated kinase activation^64^. SKS variants may disrupt this regulation, partially uncoupling mTOR activity from upstream nutrient signals. Consistent with this model, *MTOR^Δ4aa^*cells exhibit elevated basal mTOR activity but reduced dynamic responsiveness of canonical downstream outputs, alongside selective engagement of alternative signaling pathways. Altered substrate selection or RAPTOR-mediated recruitment may bias signaling away from canonical mTORC1 effectors. In line with recent studies demonstrating that mTOR regulates RNA processing and splicing independently of S6K1^53,65^, *MTOR^Δ4aa^*cells show persistent enrichment of spliceosome and RNA-processing programs across nutrient states. Transcriptomic enrichment of MYC- and E2F-driven programs likely reinforce these nuclear and gene-regulatory functions. Together, these findings support a model in which SKS mutations alter the coupling between mTOR activation and downstream outputs, favoring transcriptional and chromatin-associated regulation while attenuating canonical translational responses.

### MAPK/CaMK signaling integrates stress and transcriptional outputs in MTOR^Δ4aa^ cells

RNA-seq and proteomic analyses reveal enrichment of stress-responsive transcriptional signatures across both *MTOR^Δ4aa^*and *TSC2*^−/–^ cells, with stronger activation in *TSC2*^−/–^. These observations indicate that such signaling is a shared feature of mTOR hyperactivation^50^. In *MTOR^Δ4aa^* cells, MAPK/ERK- and Ca^2+^/CaMK-dependent pathways form a prominent regulatory axis that integrates environmental and intracellular signals with transcriptional control. These pathways are known to interface with NF-κB and other stress-responsive transcriptional programs^66–68^, but in this context appear to operate within a broader signaling architecture that prioritizes gene regulation rather than inflammatory amplification. Consistent with this, *MTOR^Δ4aa^*cells exhibit enrichment of MYC- and E2F-driven transcriptional programs alongside suppression of developmental transcription factors such as TEAD1, suggesting a shift away from lineage specification toward alternative regulatory states. Importantly, kinase activity profiling indicates reduced engagement of several canonical stress and inflammatory kinases under basal conditions in *MTOR^Δ4aa^*cells, further supporting the conclusion that inflammatory signaling is not a dominant driver of this state. Instead, *MTOR^Δ4aa^* cells appear to integrate stress and metabolic inputs through selective signaling pathways that couple environmental sensing to transcriptional regulation.

### Physiological implications and therapeutic considerations

Our study provides a systems-level framework for understanding how SKS mutations reprogram mTOR signaling networks and contribute to diverse disease phenotypes. Across datasets, mTOR gain-of-function establishes a shared baseline of nutrient-responsive and stress-associated signaling, but diverge in regulatory architecture: *MTOR^Δ4aa^* cells prioritize transcriptional and nuclear programs, whereas *TSC2^−/–^* cells engage broader proteome-level anabolic and metabolic remodeling. Rather than uniformly increasing growth signaling, *MTOR^Δ4aa^* reshapes how cells respond to nutrient cues, favoring sustained mitochondrial and gene-regulatory programs over coordinated anabolic transitions. This altered signaling profile may disrupt neuronal development, circuit function, and broader homeostatic processes, including interactions among metabolism, immunity, circadian rhythms, and sleep. Suppression of developmental transcription factors such as TEAD1 further suggests impaired lineage specification and tissue organization, providing a potential mechanistic link to neurodevelopmental phenotypes^69^.

These findings support a model in which SKS represents a signaling rewiring state, rather than simple mTOR hyperactivation. In this framework, canonical mTORC1 outputs are relatively constrained, while signaling is redistributed toward ERK/CaMK-dependent pathways and transcriptional regulation. This has important therapeutic implications: while rapamycin and rapalogs target canonical mTORC1 activity and show variable benefit in SKS, our data suggest that combination strategies targeting downstream rewired pathways, particularly MAPK/ERK and CaMK-dependent signaling, may be required to restore balanced signaling outputs and improve therapeutic precision.

In summary, *MTOR^Δ4aa^* cells adopt a non-canonical nutrient-response state distinct from *TSC2* loss. Despite shared increases in mitochondrial programs, *MTOR^Δ4aa^* cells exhibit constrained proteomic dynamics, selective engagement of MAPK/ERK- and CaMK-driven signaling pathways, and redistribution of mTOR outputs toward transcriptional and gene-regulatory programs. These findings support a model in which SKS represents a distinct mode of mTOR dysregulation defined by signaling rewiring rather than uniform canonical pathway activation, with implications for mechanism-based therapeutic strategies.

### Limitations of the study

This study has several limitations. Experiments were performed in human U2OS cells, which do not capture tissue-specific, developmental, or circuit level contexts of mTOR signaling^7^. In addition, we examined only a single SKS allele, *MTOR^(ΔR1480-C1483)^* and future studies should test additional variants and extend these findings to disease-relevant cell types, including neuronal and glial systems, as well as in animal models, to determine which aspects of this signaling architecture generalize across SKS mutations. Integration with *in vivo* pathophysiology, including behavioral and metabolic phenotypes, will be essential to establish translational relevance.

## Resource availability

### Lead contact

Further information and requests for resources and reagents should be directed to and will be fulfilled by the lead contact, Andrew C. Liu

### Materials availability

All materials and reagents used in this study are available from the lead contact upon request.

### Data and code availability

- All data needed to evaluate the conclusions in the paper are present in the paper and/or Supplementary Materials.
- Raw proteomics and phosphoproteomics datasets have been deposited in the MassIVE repository, with the identifier MSV000101560
- RNA-seq data have been deposited in the Gene Expression Omnibus (GEO) database under the accession code GSE330231
- Python scripts have been deposited to GitHub: https://github.com/Mrbland/SKS_phospho_processing_scripts

## Acknowledgments

We would like to thank the UF ICBR NextGen sequencing core facility for RNA sequencing support. Research reported in this work was supported by NIH NINDS R01 NS117457 to ACL, NIH NINDS F31 F31NS147396, NIH T32 GM 007377, and NIH T32 GM 153586 to CRC. Research in the laboratory of JCC is supported by NIH grants R01 DK124068 and R21 AG082480. The content is solely the responsibility of the authors and does not necessarily represent the official views of the NIH. Cartoons in figures were created with BioRender.com, license to JCC.

## Author contributions

ACL, JCC, and CRC conceptualized and designed the study. ACL, CRC, and JCC acquired the funding. ACL, JCC, and JM coordinated and supervised the investigation. CRC, YS, HH, EKG, CH, and JM performed research and analyzed data. CRC, ACL, JCC contributed to critical interpretation of the data. CRC drafted the manuscript with input from ACL and JCC. All authors edited the manuscript.

## Declaration of Interests

The authors declare no competing interests.

## Supplemental Figure Legends

**Supplemental Figure 1.**
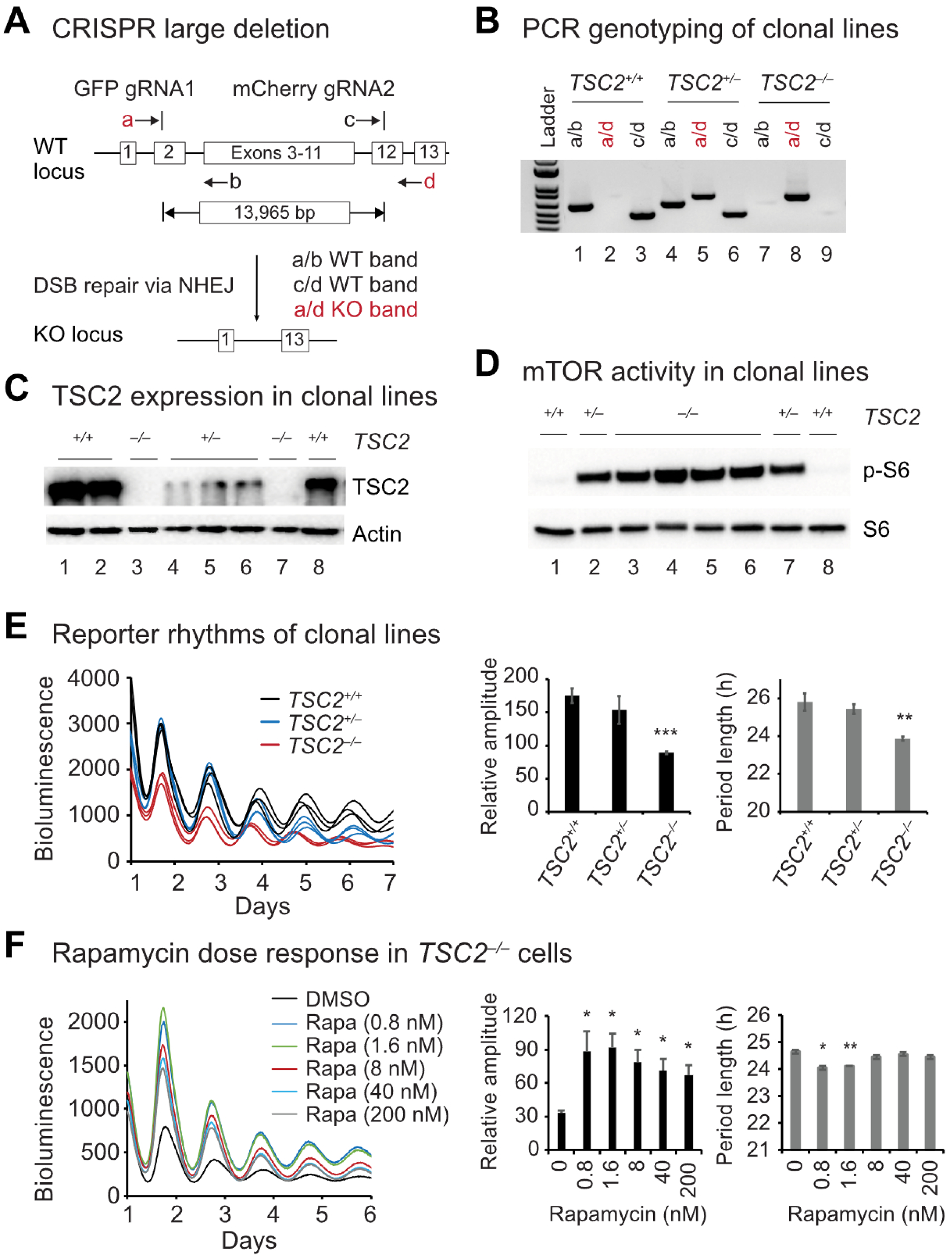
Generation and validation of *TSC2* knockout human U2OS cells. (A) Schematic of CRISPR–Cas9–mediated deletion of the human *TSC2* locus using two sgRNAs flanking exons 2–12, generating a large (∼13.9 kb) genomic deletion. Primer locations and expected PCR products for wild-type and deletion alleles are shown. (B) PCR genotyping of clonal cell lines using multiple primer pairs distinguishes wild-type (WT), heterozygous, and homozygous *TSC2* knockout clones. The F51/R32 primer pair yields a ∼662 bp product in *TSC2^−/−^* clones, whereas the wild-type allele does not amplify under standard PCR conditions. (C) Immunoblot analysis of TSC2 protein expression in representative clonal lines confirms loss of TSC2 in homozygous knockout cells. (D) Immunoblot analysis of mTORC1 activity showing increased phosphorylation of ribosomal protein S6 (p-S6 S235/236) in *TSC2^−/−^*cells compared with WT and heterozygous controls. (E) Representative circadian bioluminescence rhythms (left) and quantification (right) showing shortened period length and reduced rhythm amplitude in *TSC2^−/−^* cells relative to WT and heterozygous clones. (F) Rapamycin treatment partially restores circadian rhythm amplitude in *TSC2^−/−^* cells in a dose-dependent manner, with representative traces (left) and amplitude quantification (right). Data are shown as mean ± SD. Statistical significance was determined using Student’s t test: *p < 0.05, **p < 0.01, compared with controls.

**Supplemental Figure 2.**
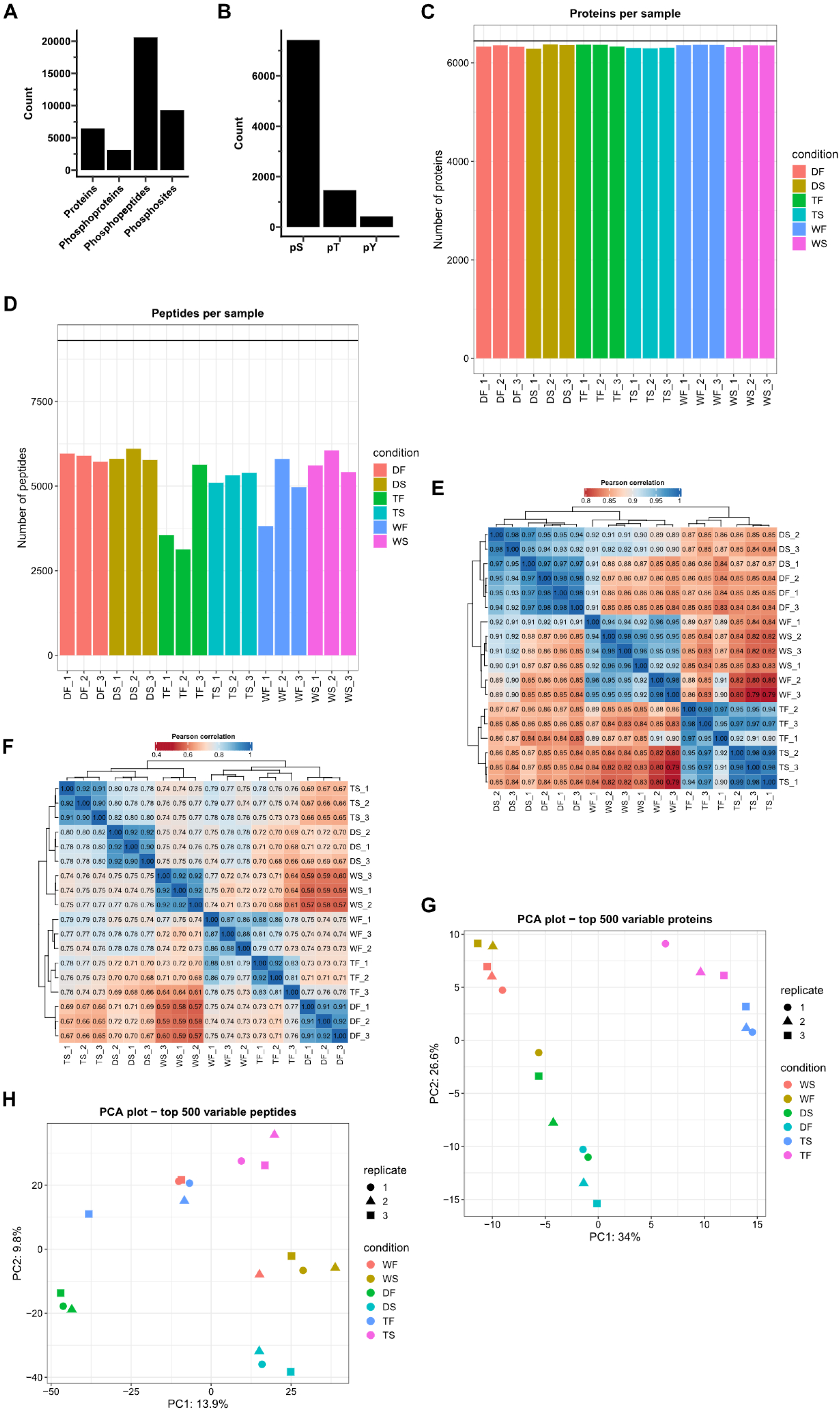
Omics data quality and clustering. (A) Bar plot showing the number of detected proteins, phosphoproteins, phosphopeptides, and unique phosphosites that passed quality control filtering. (B) Bar plot showing the number of phosphorylated amino acids detected in the phosphoproteomics dataset (pS = phosphor-Serine; pT = phosphor-Threonine; pY = phospho-Tyrosine). (C-D) Number of (C) proteins or (D) phosphopeptides detected in each condition and replicate (Δ4aa = *MTOR^(ΔR1480-C1483)^*; TSC = *TSC2^−/−^*; WT = wild-type). (E-F) Pearson correlation heatmaps for (E) total proteins and (F) total phosphopeptides (F). Samples are hierarchically clustered based on pairwise Pearson correlation (distance = 1 – r), with dendrograms indicating sample similarity. Correlation strength is color coded (blue = highly correlated, red = weakly correlated). (G-H) Principal component analysis (PCA) plots of the top 500 variable proteins (G) and phosphopeptides (H). Each point represents a biological replicate, colored by conditions (WF = wild-type glucose-fed; WS = wild-type glucose-depleted; DF = *MTOR^Δ4aa^* glucose-fed; DS = *MTOR^Δ4aa^* glucose-depleted; TF = *TSC2*^−/–^ glucose-fed; TS = *TSC2*^−/–^ glucose-depleted) and shaped by replicate number (n=3 per condition).

**Supplemental Figure 3.**
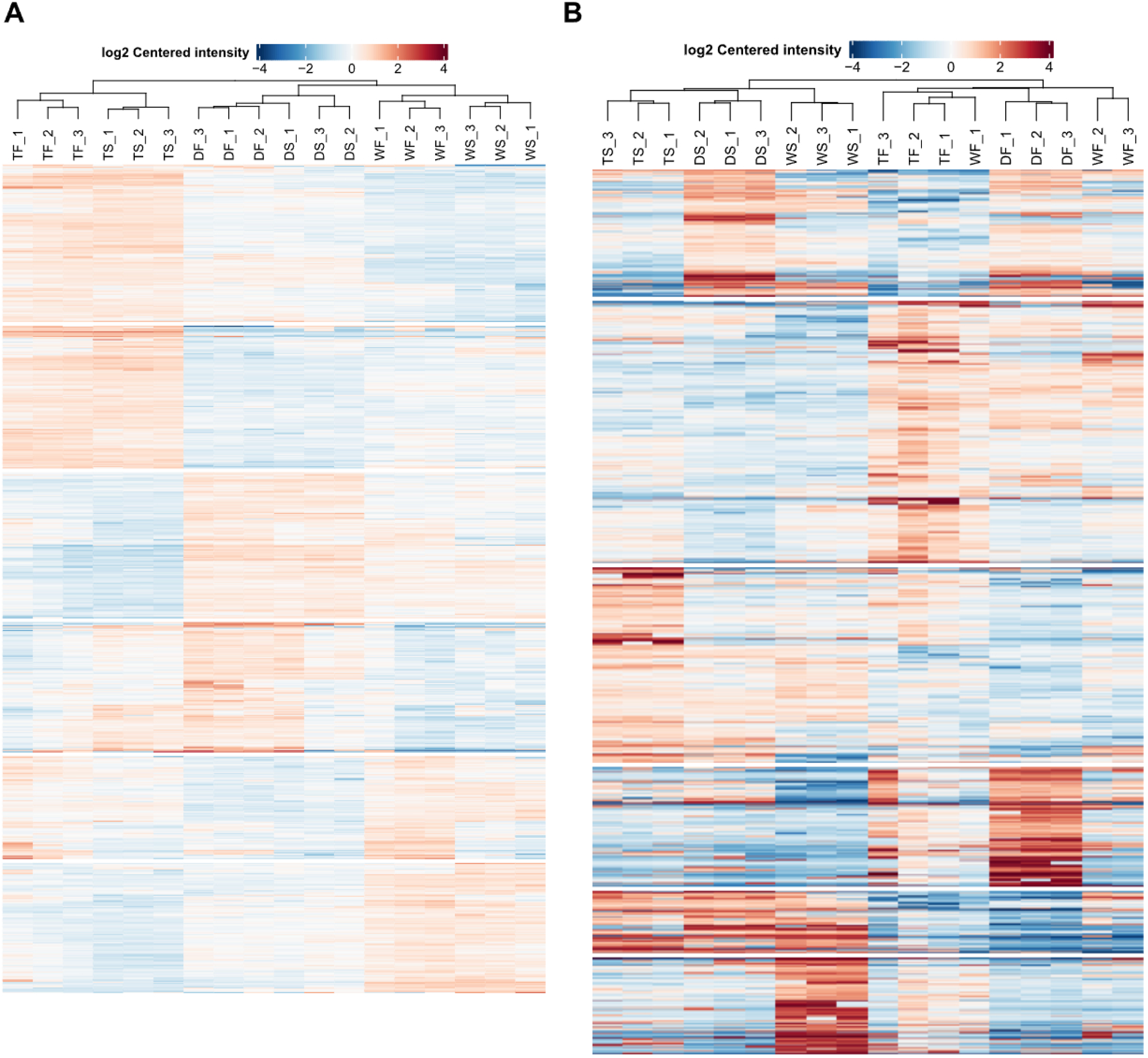
Hierarchical clustering of total protein and phosphosite intensity profiles. (A) Heatmap of Log_2_-centered intensities for total protein abundance across all samples. Rows represent proteins and columns represent samples. Intensities were mean-centered per protein, and both rows and columns were hierarchically clustered using Pearson correlation distance (1 – r) and average linkage. Dendrograms reflect sample similarity (top) and protein co-expression patterns (left). (B) Heatmap of Log_2_-centered phosphosite intensities. Phosphosites are mean-centered across samples prior to clustering as in (A). Rows represent individual phosphosites, and columns represent biological replicates across genotype and nutrient conditions. Color scales represent relative abundance (mean-centered values): red = higher than mean, blue = lower than mean. (Δ_F# = *MTOR^Δ4aa^* glucose-fed; Δ_S# = *MTOR^Δ4aa^* glucose-depleted; T_F# = *TSC2*^−/–^ glucose-fed; T_S# = *TSC2*^−/–^glucose-depleted; W_F# = wild-type glucose-fed; W_S# = wild-type glucose-depleted)

**Supplemental Figure 4.**
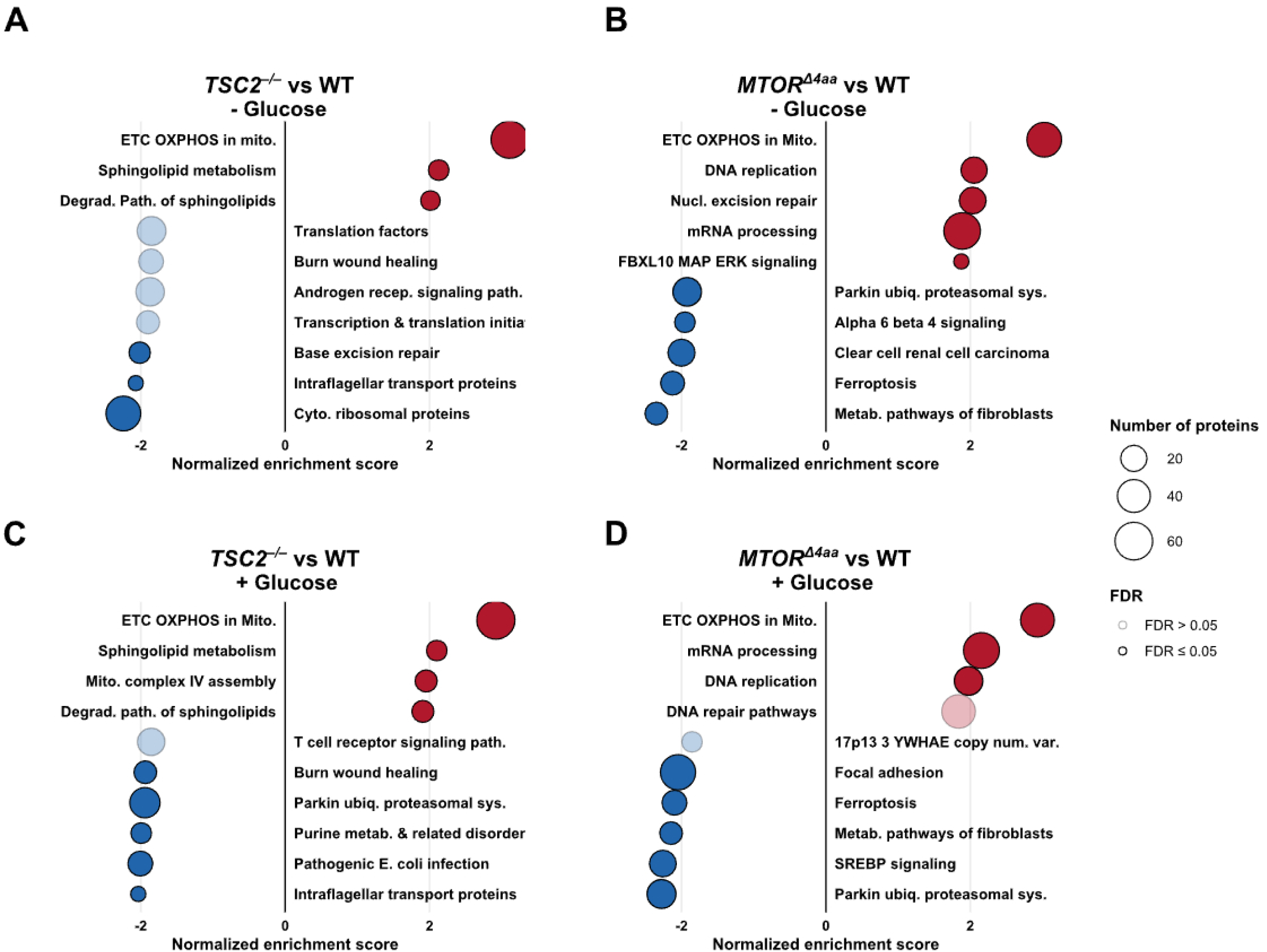
WikiPathway gene set enrichment comparisons of *MTOR^Δ4aa^* and *TSC2^−/–^*cells against WT cells under glucose-depleted and glucose-fed conditions. (A-D) Gene Set Enrichment Analysis (GSEA) using the WikiPathways database comparing. (A) *TSC2*^−/–^ against WT cells under glucose-depleted conditions, (B) *MTOR^Δ4aa^* against WT cells under glucose-depleted conditions, (C) *TSC2*^−/–^ against WT cells under glucose-fed conditions, and (D) *MTOR^Δ4aa^* against WT cells under glucose-fed conditions. Each plot displays the top 10 enriched pathways or complexes ranked by normalized enrichment score derived from Log2 Fold Change values. Bubbles are sized by the number of proteins mapped to each gene set, colored by direction of enrichment (red = upregulated, blue = downregulated), and shaded by False Discovery Rate (solid bubbles = FDR ≤ 0.05; transparent bubbles = FDR > 0.05).

**Supplemental Figure 5.**
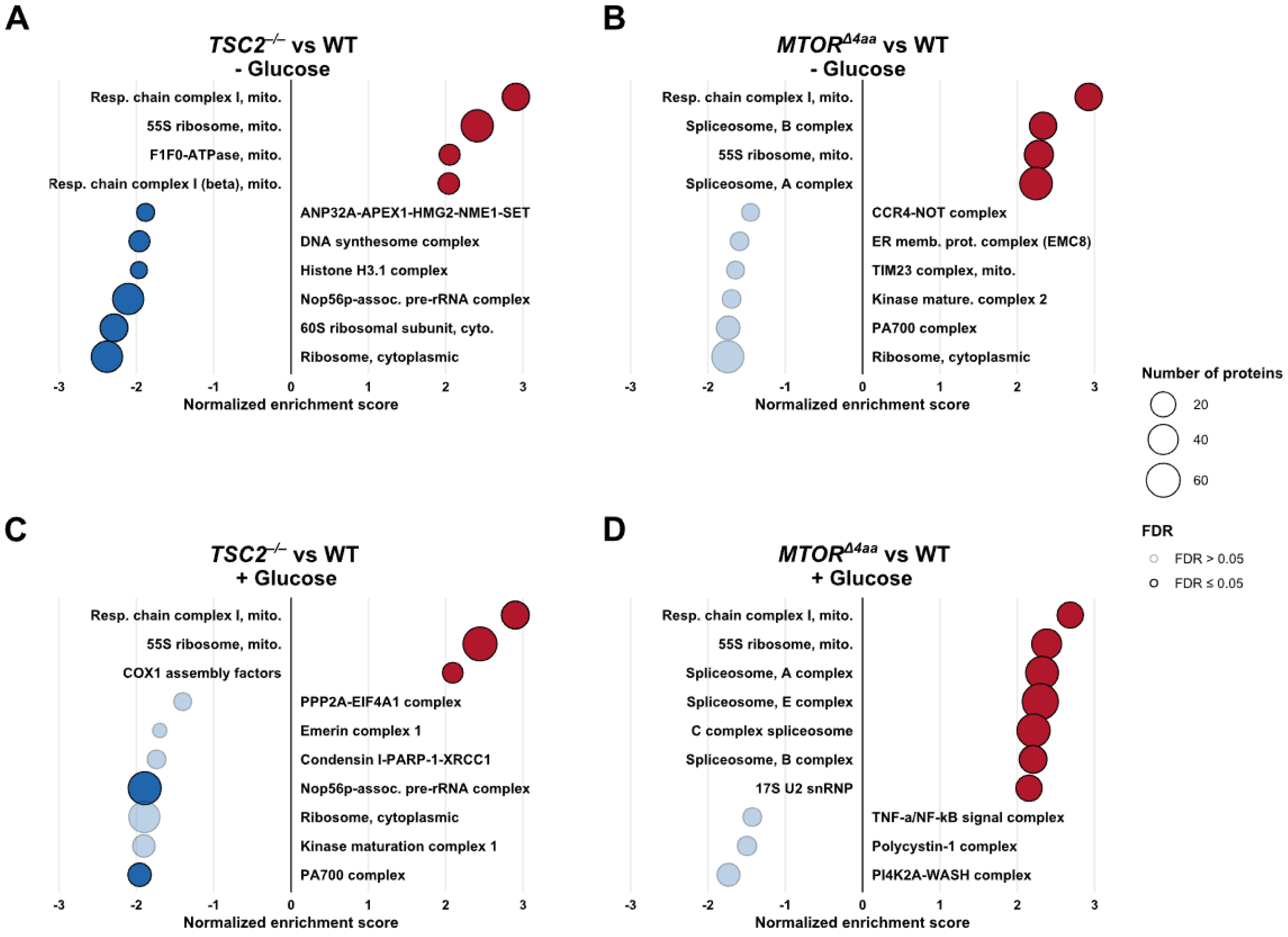
CORUM protein complex enrichment comparisons of *MTOR^Δ4aa^* and *TSC2^−/–^* cells against WT cells under glucose-depleted and glucose-fed conditions. Gene Set Enrichment Analysis (GSEA) using the CORUM protein complex database comparing. (A) glucose-depleted *TSC2*^−/–^ cells against glucose-depleted WT cells, (B) glucose-depleted *MTOR^Δ4aa^* cells against glucose-depleted WT cells, (C) glucose-fed *TSC2*^−/–^ cells against glucose-fed WT cells, and (D) glucose-fed *MTOR^Δ4aa^* cells against glucose-fed WT cells. Each plot displays the top 10 enriched pathways or complexes ranked by normalized enrichment score derived from Log_2_ Fold Change values. Bubbles are sized by the number of proteins mapped to each gene set, colored by direction of enrichment (red = upregulated, blue = downregulated), and shaded by False Discovery Rate (solid bubbles = FDR ≤ 0.05; transparent bubbles = FDR > 0.05).

**Supplemental Figure 6.**
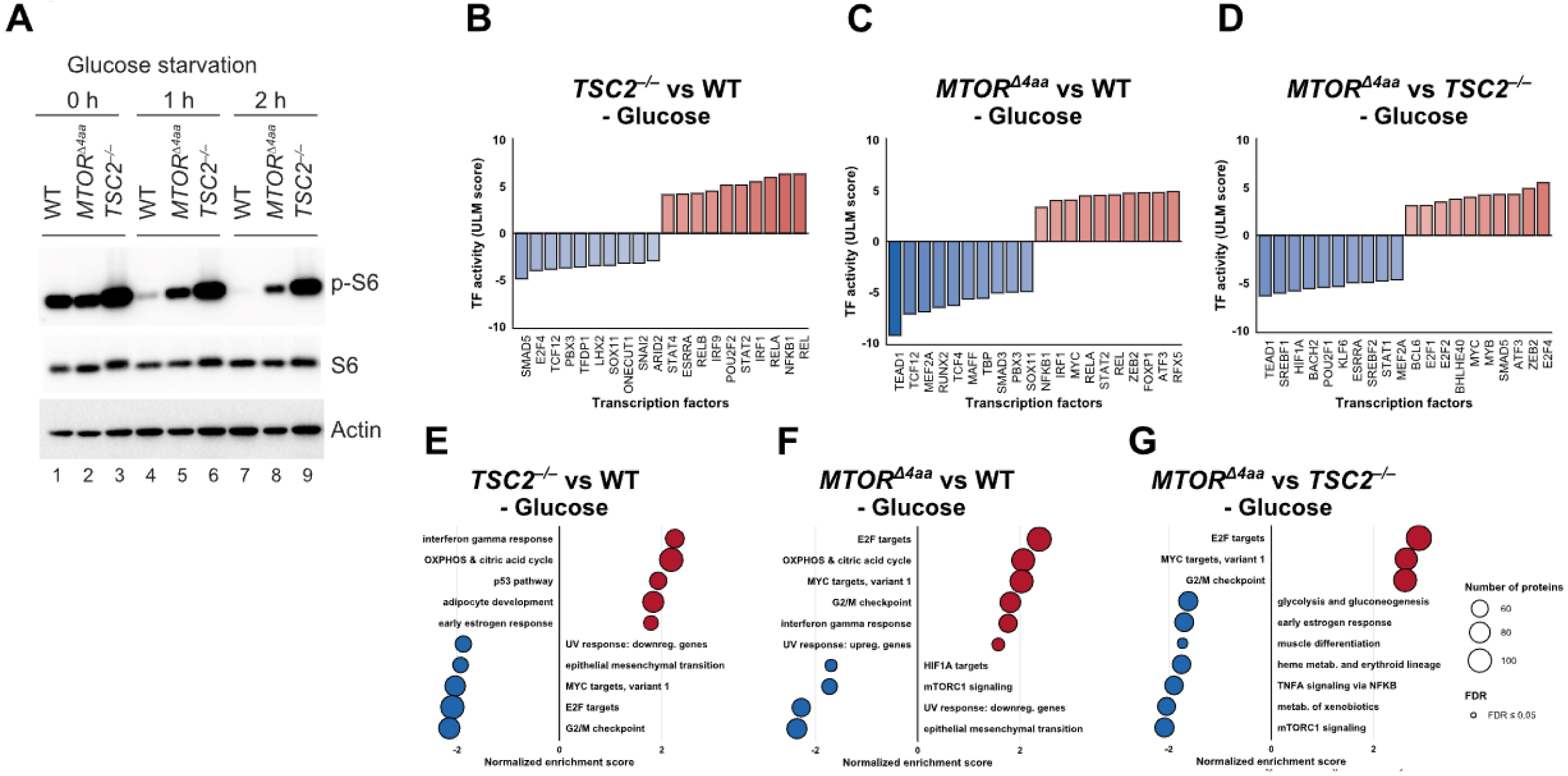
Distinct basal transcriptional states in *MTOR^Δ4aa^* and *TSC2^−/–^* cells under glucose depletion. **(A)** WT, *MTOR^Δ4aa^*, and *TSC2^−/−^* U2OS cells were subjected to glucose depletion for 0 h, 1 h, or 2 h. Representative immunoblots show phosphorylation of ribosomal protein S6 (p-S6), total S6, and actin as a loading control. Lanes 1–3: 0 h; lanes 4–6: 1 h; lanes 7–9: 2 h. *TSC2^−/−^*cells maintain elevated mTOR activity during prolonged starvation, whereas *MTOR^Δ4aa^*cells exhibit intermediate suppression relative to WT. Samples from 2 h glucose depletion (lanes 7–9) were used for RNA-seq. (B-D) Inferred differential transcription factor activities across pairwise comparisons. Bar plots display the top 10 positively and negatively regulated transcription factors ranked by ULM score, with red indicating relatively higher inferred activity and blue indicating relatively lower inferred activity for *TSC2^−/−^* vs WT (B), *MTOR^Δ4aa^* vs WT (C), and *TSC2^−/−^* vs *MTOR^Δ4aa^* (D). (E-G) GSEA using the MSigDB Hallmark Gene Set database for *TSC2^−/−^* vs WT (E), *MTOR^Δ4aa^* vs WT (F), and *TSC2^−/−^* vs *MTOR^Δ4aa^* (G). Each plot displays the top 10 enriched gene sets ranked by normalized enrichment score derived from Log_2_ Fold Change values. Bubbles are sized by the number of proteins mapped to each gene set, colored by direction of enrichment (red = upregulated, blue = downregulated), and shaded by False Discovery Rate (solid bubbles = FDR ≤ 0.05; transparent bubbles = FDR > 0.05).

## Supplemental Table Legends

**Table S1. Full DEP2 proteomics output, related to Figures 1-4 and S1-5.** Complete DEP2 differential protein abundance output from quantitative proteomics analysis of WT, *MTOR^Δ4aa^*, and *TSC2^−/−^*cells under glucose-depleted and glucose-refed conditions. The table includes protein identifiers, gene annotations, normalized abundance values, pairwise comparison results, log2 fold changes, statistical outputs, and adjusted p-values used for downstream proteomic and enrichment analyses.

**Table S2. Full DEP2 phosphoproteomics output, related to Figures 5-6 and S2.**

Complete DEP2 differential phosphorylation output from phosphoproteomics analysis of WT, *MTOR^Δ4aa^*, and *TSC2^−/−^*cells under glucose-depleted and glucose-fed conditions. The table includes phosphosite identifiers, parent protein and gene annotations, normalized and protein-corrected phosphosite abundance values, pairwise comparison results, log2 fold changes, statistical outputs, and adjusted p-values used for downstream kinase enrichment analyses.

**Table S3. GO Biological Process overrepresentation analysis of *MTOR^Δ4aa^* vs WT and *TSC2^−/−^*vs WT comparisons, related to Figures 2-3.**

Gene Ontology Biological Process overrepresentation analysis results for shared and genotype-specific differentially abundant proteins identified in *MTOR^Δ4aa^* vs WT and *TSC2^−/−^* vs WT comparison under glucose-depleted and glucose-refed conditions. The table includes enriched biological processes, enrichment statistics, adjusted p-values, associated gene or protein sets, and comparison-specific annotations.

**Table S4. WikiPathway gene set enrichment analysis of *MTOR^Δ4aa^* vs WT and *TSC2^−/−^* vs WT comparisons, related to Figure S4.**

Gene set enrichment analysis results using the WikiPathways database for *MTOR^Δ4aa^* vs WT and *TSC2^−/−^* vs WT proteomic comparisons under glucose-depleted and glucose-refed conditions. The table includes enriched pathways, enrichment statistics, normalized enrichment scores, false discovery rates, associated leading-edge or core-enrichment proteins, and comparison-specific annotations.

**Table S5. CORUM protein complex gene set enrichment analysis of *MTOR^Δ4aa^*vs WT and *TSC2^−/−^*vs WT comparisons, related to Figure S5.**

Gene set enrichment analysis results using the CORUM protein-complex database for *MTOR^Δ4aa^* vs WT and *TSC2^−/−^* vs WT proteomic comparisons under glucose-depleted and glucose-refed conditions. The table includes enriched protein complexes, enrichment statistics, normalized enrichment scores, false discovery rates, associated leading-edge or core-enrichment proteins, and comparison-specific annotations.

**Table S6. Hallmark50 gene set enrichment analysis of intragenotype glucose-refed vs glucose-depleted comparisons, related to Figure 4.**

Gene set enrichment analysis results using the MSigDB Hallmark50 database for glucose-refed vs glucose-depleted proteomic comparisons within WT, *MTOR^Δ4aa^*, and *TSC2^−/−^* cells. The table includes enriched Hallmark gene sets, enrichment statistics, normalized enrichment scores, false discovery rates, associated leading-edge or core-enrichment proteins, and comparison-specific annotations.

**Table S7. Full DEP2 RNA-seq output, related to Figure S6.**

Complete DEP2 differential gene expression output from RNA-seq analysis of WT, *MTOR^Δ4aa^*, and *TSC2^−/−^* cells under glucose-depleted conditions. The table includes gene identifiers, gene annotations, normalized expression values, pairwise comparison results, log2 fold changes, statistical outputs, and adjusted p-values used for downstream transcriptome-level analyses.

**Table S8. Inferred transcription factor activity analysis output, related to Figure S6.**

Complete inferred transcription factor activity analysis output generated from RNA-seq differential expression results using DoRothEA and decoupleR. The table included inferred transcription factors, statistical test, comparison-specific annotations, enrichment scores, and p-values.

**Table S9. Hallmark50 gene set enrichment analysis of differentially expressed genes from RNA-seq, related to Figure S6.**

Gene set enrichment analysis results using the MSigDB Hallmark50 database for RNA-seq-derived differential expression comparisons among WT, *MTOR^Δ4aa^*, and *TSC2^−/−^* cells under glucose-depleted conditions. The table includes enriched Hallmark gene sets, enrichment statistics, normalized enrichment scores, false discovery rates, associated leading-edge or core-enrichment genes, and comparison-specific annotations.

**Table S10. Serine/threonine kinase enrichment output from The Kinase Library, related to Figures 5-6 and Table S2.**

Complete serine/threonine kinase enrichment results generated from phosphoproteomics data using The Kinase Library. The table includes predicted upstream serine/threonine kinases, kinase gene names, kinase families, enrichment scores, p-values, adjusted p-values, number of phosphosites and their direction of regulation, and specific comparison annotations.

**Table S11. Tyrosine kinase enrichment output from The Kinase Library, related to Figures 5-6 and Table S2.**

Complete tyrosine kinase enrichment results generated from phosphoproteomics data using The Kinase Library. The table includes predicted upstream serine/threonine kinases, kinase gene names, kinase families, enrichment scores, p-values, adjusted p-values, number of phosphosites and their direction of regulation, and specific comparison annotations.

## Notes

### Competing Interest Statement

The authors have declared no competing interest.

